# Gut microbiota and age shape susceptibility to clostridial enteritis in lorikeets under human care

**DOI:** 10.1101/2021.09.08.459483

**Authors:** David Minich, Christopher Madden, Mauricio A. Navarro, Leo Glowacki, Kristen French-Kim, Willow Chan, Morgan V. Evans, Kilmer Soares, Ryan Mrofchak, Rushil Madan, Gregory A. Ballash, Krista LaPerle, Subhadeep Paul, Yael Vodovotz, Francisco A. Uzal, Margaret Martinez, Jennifer Hausmann, Randall E. Junge, Vanessa L. Hale

## Abstract

**Background:** Enteritis is a common cause of morbidity and mortality in lorikeets that can be challenging to diagnose and treat. In this study, we examine gut microbiota in two lorikeet flocks with enteritis (Columbus Zoo and Aquarium – CZA; Denver Zoo - DZ). Since 2012, the CZA flock has experienced repeated outbreaks of enteritis despite extensive diet, husbandry, and clinical modifications. In 2018, both CZA and DZ observed a spike in enteritis. Recent research has revealed that the gut microbiota can influence susceptibility to enteropathogens. We hypothesized that a dysbiosis, or alteration in the gut microbial community, was making some lorikeets more susceptible to enteritis, and our goal was to characterize this dysbiosis and determine the features that predicted susceptibility.

**Results:** We employed 16S rRNA sequencing to characterize the cloacal microbiota in lorikeets (CZA n = 67, DZ n = 24) over time. We compared the microbiota of healthy lorikeets, to lorikeets with enteritis, and lorikeets susceptible to enteritis, with “susceptible” being defined as healthy birds that subsequently developed enteritis. Based on sequencing data, culture, and toxin gene detection in intestinal contents, we identified *Clostridium perfringens* type A (CZA and DZ) and *C. colinum* (CZA only) at increased relative abundances in birds with enteritis. Histopathology and immunohistochemistry further identified the presence of gram-positive bacilli and *C. perfringens,* respectively, in the necrotizing intestinal lesions. Finally, using Random Forests and LASSO models, we identified several features (young age and the presence of *Rhodococcus fascians* and *Pseudomonas umsongensis*) associated with susceptibility to clostridial enteritis.

**Conclusions:** We identified *C. perfringens* type A and *C. colinum* associated with lorikeet necrohemorrhagic enteritis at CZA and DZ. Susceptibility testing of isolates lead to an updated clinical treatment plan which ultimately resolved the outbreaks at both institutions. This work provides a foundation for understanding gut microbiota features that are permissive to clostridial colonization and host factors (e.g. age, prior infection) that shape responses to infection.

## Background

Enteritis is one of the most common causes of morbidity and mortality in captive and free-living lorikeets and lories, and outbreaks have been reported at multiple zoological institutions [1–5]. In an informal survey of 12 North American zoos that house 10 or more lorikeets, 11 of the 12 zoos reported a history of enteritis in their flocks, and 50% of the zoos reported recurrent outbreaks of enteritis (Junge, Hausmann, unpublished data). In at least five of these zoos, repeated cultures failed to identify an etiologic agent, and a combination of broad-spectrum antimicrobials were employed as empiric therapy. This approach increases the risk of promoting antimicrobial resistance [6]; moreover, in many cases, it also failed to resolve the outbreak or prevent recurrences of enteritis.

The microbiome is a collection of bacteria, Archaea, viruses, and microbial eukaryotes that live on or in hosts such as lorikeets. The microbiome is increasingly being realized as a source for biomarkers that predict disease and clinical outcomes [7–9] and serve as targets for therapeutics [10] (e.g. probiotics, prebiotics, phage therapy, dietary modification, microbiota transplants). Over the past decade, studies on human and animal microbiomes have increased exponentially [11], and we have learned that these microbial communities are critical to host health. The gut microbiome, for example, plays an important role in nutrient acquisition and metabolism [12], immune development [13], and pathogen defense [14, 15]. There are a growing number of studies on avian [16–18] and wildlife microbiota [19–21], and these studies are providing key insights into health and disease using minimally or non-invasive sampling techniques [22–25]. Microbiome studies are also being used to inform conservation efforts in wildlife species [26–28]. Recent research has further revealed that the gut microbiota influences susceptibility to enteropathogens. For example, *Clostridium*, *Campylobacter*, and *Salmonella* species, all of which are common agents in avian enteritis [1, 2, 5, 29–34], are adept at invading dysbiotic (or already disrupted) microbial communities but not at colonizing healthy microbial communities [35–37].

In this study, we examined the microbiota over time in two lorikeet flocks (Columbus Zoo and Aquarium [CZA], Denver Zoo [DZ]) that experienced one or more outbreaks of enteritis. Between 2012 and 2018, the CZA lorikeet flock experienced repeated outbreaks of enteritis despite extensive efforts to resolve these issues through nutrition, sanitation, medication, and habitat and husbandry modifications [38]. In 2018, lorikeet morbidity and mortality events spiked, and necropsy reports consistently identified severe and necrotizing enteritis in these lorikeets, but bacterial cultures were frequently negative or variable (e.g. *Escherichia coli, Enterococcus spp.*). Around the same time, DZ was also managing an outbreak of enteritis in their lorikeet flock. We took a novel approach to lorikeet enteritis and sampled healthy and sick birds at CZA and DZ over 10 months. We hypothesized that a dysbiosis, or alteration in the gut microbial community, was making some lorikeets more susceptible to enteritis. Hence, our goal was to characterize this dysbiosis and determine if and what features predicted susceptibility.

## Results

### Columbus Zoo and Aquarium lorikeet enteritis

We sampled a total of 67 lorikeets at the CZA between November 2018 and September 2019. During this time, 34 lorikeets developed enteritis one or more times while the remaining birds (n = 33) never developed enteritis (Table 1). Birds with enteritis were identified through clinical signs (diarrhea, lethargic, fluffed). “Enteritis” samples collected from the same bird within two weeks of the initial enteritis sample were considered a single case. Enteritis samples collected beyond 2 weeks from the initial sample in the same bird were counted as a second case of enteritis. There were no significant differences by sex in the number of lorikeets that did or did not develop enteritis (χ^2^ = 5.7, *p =* 0.06; χ^2^ between males and females only, excluding “unknown” = 0.12, *p* = 0.73). However, there was a significant difference by age (χ^2^ = 9.7, *p =* 0.02) and by species (χ^2^ with all species = 7.2, *p* = 0.07; χ^2^ between rainbow and coconut lorikeets only (the two dominant species in this flock) = 6, *p* = 0.01), with enteritis occurring more in younger birds (< 2 years old) and in coconut lorikeets. Birds that survived a previous episode of enteritis also had a 2.2 times increased risk of developing future enteritis; although, this was only marginally significant [95% CI: 0.99-5.29; p=0.051]. We also observed an increase in enteritis cases between December 2018 and March 2019 (Fig. 1a).

**Figure 1.**
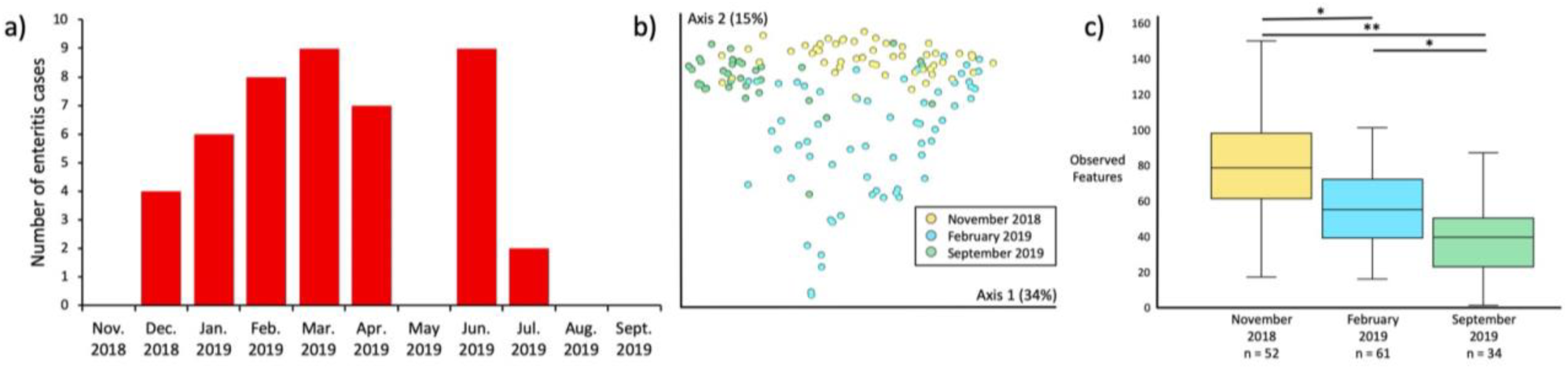
Seasonality in enteritis and gut microbiota in healthy lorikeets. **a)** Number of enteritis cases by month in Columbus Zoo and Aquarium lorikeets. Enteritis samples collected from the same bird within two weeks of the initial enteritis sample were considered a single case. Enteritis samples collected beyond 2 weeks from the initial sample in the same bird were counted as a second case of enteritis. Columbus Zoo & Aquarium healthy birds only: **b)** Microbial community composition (Weighted UniFrac) and **c)** diversity (Observed Features) by season (November 2018, February 2019, September 2019). There were significant shifts in microbial composition by season (PERMANOVA *p* = 0.001), and microbial diversity decreased significantly between November 2018 and September 2019 (Kruskal-Wallis \**p* < 0.001, \*\**p* < 0.0001). (Also see **Additional File 6.**)

**Table 1.** Columbus Zoo and Aquarium (CZA) Lorikeet Demographics. Number and percent of Columbus Zoo & Aquarium lorikeets that ever or never had enteritis between Nov. 2018 and Sept. 2019 by sex, age, and lorikeet species.

|  | Lorikeets that had<br>**one or more**<br>**episodes of**<br>**enteritis** between<br>Nov. 2018-Sept.<br>2019 | Lorikeets that<br>**never had enteritis**<br>between<br>Nov. 2018-Sept.<br>2019 | *p*-value |
| --- | --- | --- | --- |
| **Sex (n, %)** |  |  |  |
| Male | 9 (13.4 %) | 14 (20.9 %) | p = 0.06 (all)<br>p = 0.73 (male<br>vs. female<br>only) |
| Female | 11 (16.4 %) | 14 (20.9 %) |  |
| Unknown | 14 (20.9 %) | 5 (7.5 %) |  |
| **Age (n, %)** |  |  |  |
| 0-2 years | 19 (28.3%) | 7 (10.4%) | p = 0.02 |
| 3-5 years | 7 (10.4%) | 16 (23.9 %) |  |
| 6-10 years | 6 (9.0%) | 6 (9.0%) |  |
| >11 years | 2 (3.0 %) | 4 (6.0 %) |  |
| **Species (n, %)** |  |  |  |
| Rainbow | 8 (11.9 %) | 17 (25.4 %) | p = 0.07 (all)<br>p = 0.01<br>(rainbow vs.<br>coconut only) |
| Coconut | 23 (34.3 %) | 13 (19.4 %) |  |
| Marigold | 1 (1.5 %) | 0 (0 %) |  |
| Lory | 2 (3 %) | 3 (4.5 %) |  |

### Gut microbiota by demographics and season

We did not observe significant differences in microbial composition or diversity (Shannon, Observed Features) by sex or species; although, by age, some differences in composition were detected via the Unweighted, but not the Weighted UniFrac metric (**Additional File 4**, alpha-diversity, all Kruskal-Wallis *p* > 0.05; **Additional File 5**). There were also significant differences in microbial composition and diversity by season (November 2018, February 2019, September 2019) with diversity declining significantly over time (Fig. 1b,c, Weighted UniFrac, PERMANOVA *p* = 0.001; Observed Features Kruskal-Wallis, all *p* < 0.001, **Additional File 6**).

### Gut microbiota changes with enteritis

As compared to healthy lorikeets, we observed a significant decrease in microbial diversity and altered microbial composition in lorikeets with enteritis or lorikeets that died or were euthanized as a result of enteritis (denoted “post-mortem” lorikeets) (Fig. 2, Observed Features Kruskal-Wallis all *p* < 0.0005, Weighted UniFrac PERMANOVA *p* = 0.001; **Additional File 7**). The top differentially abundant microbe between healthy birds and birds with enteritis was *Clostridium perfringens*, which was significantly increased in lorikeets with enteritis (Fig. 2c; ANCOM W=1098, no post-mortem birds included in this analysis). There were three other differentially abundant clostridia, including *C. colinum, C. neonatale,* and *C. paraputrificum* which were also all increased in birds with enteritis (**Additional File 8**). We then performed this analysis on a subset of birds (n = 25) for which we had matched samples at both healthy and enteritis time points and we again observed a significantly increased abundance of clostridia including *C. perfringens* and *C. neonatale* in these birds when they had enteritis (Fig. 2c, ANCOM W = 788, **Additional File 9**).

**Figure 2.**
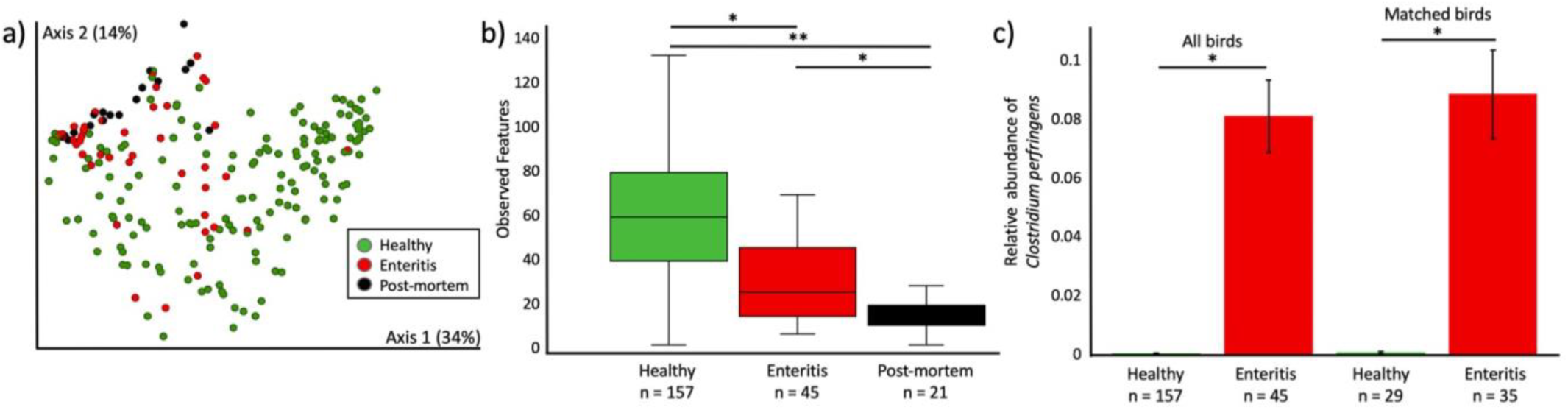
Microbial community analysis in CZA lorikeets with enteritis. Microbial composition and diversity in healthy lorikeets, lorikeets with enteritis, and lorikeets that died or were euthanized due to enteritis (postmortem). **a)** Microbial composition (Weighted UniFrac) was significantly altered (PERMANOVA *p* = 0.001) and **b)** microbial diversity (Observed Features) was significantly decreased (Kruskal-Wallis \**p* < 0.0005, \*\**p* < 1 x 10^-9^) in lorikeets with enteritis or postmortem lorikeets. (Also see **Additional File 7**.) **c)** The relative abundance of *Clostridium perfringens* was significantly increased lorikeets with enteritis across all birds (ANCOM, W = 1098) and across matched birds (ANCOM, W = 788). “All birds” included samples from healthy birds that never got enteritis. “Matched birds” included only 25 birds that had both enteritis and healthy samples.

### Culture, genotyping, and susceptibility testing

After identifying clostridial DNA in our sequencing data, we employed anaerobic culture of lorikeet intestinal contents to look for the presence of viable clostridia in the gut. Intestinal content was collected from a total of 13 lorikeets that died or were euthanized due to enteritis. Contents were cultured on TSA agar with 5% sheep blood. All colonies with unique morphology were picked between 24 - 48 hours and underwent MALDI-TOF identification. We identified clostridia in 7 out of the 13 intestinal content samples including *C. perfringens* in 6 of these 7 samples (Table 2). Other microbes that were identified in culture included: *Escherichia coli* (in 3 out of 13 samples), and *C. paraperfringens* or *C. baratii*, *C. paraputrificum,* and *C. disporicum* each in 1 out of 13 samples. We did not culture *C. colinum* in any sample.

**Table 2.** Intestinal Content Isolates from CZA Lorikeets.

| Bird ID | Intestinal contents, growth in culture? | MALDI identification | *cpa* (alpha toxin) | *cpb2* (beta2 toxin) | Inferred *C. perfringens* toxinotype | Selected for susceptibility testing? |
| --- | --- | --- | --- | --- | --- | --- |
| 104073 | No | None |  |  |  |  |
| 113096 | Yes | *Clostridium perfringens* and *Clostridium disporicum* | + | + | A |  |
| 115057 | No | None |  |  |  |  |
| 115069 | Yes | *Enterococcus faecium* |  |  |  |  |
| 115094 | No | None |  |  |  |  |
| 116046 | Yes | *Escherichia coli* |  |  |  |  |
| 118064 | Yes | *Clostridium perfringens* | + | + | A |  |
| 118071 | Yes | *Clostridium perfringens*, *Escherichia coli*, and *Clostridium paraperfringens* or *Clostridium baratii* | + | + | A | yes |
| 118075 | Yes | *Clostridium perfringens* | + | + | A | yes |
| 118089 | Yes | *Clostridium perfringens*, *Escherichia coli*, an organism with no ID | + | + | A |  |
| 118115 | No | None |  |  |  |  |
| 118118 | Yes | *Clostridium perfringens* and *Clostridium disporicum* | + | - | A |  |
| 118119 | Yes | *Clostridium paraputrificum* |  |  |  |  |

We then performed toxinotyping on the 6 *C. perfringens* isolates. All isolates contained the *cpa* gene (encoding alpha toxin) and were identified as *C. perfringens* type A. Five out of 6 isolates also contained the *cpb2* gene (encoding beta 2 toxin) (Table 2). Susceptibility testing was performed on 2 isolates. Both isolates were susceptible to metronidazole and penicillin, and one was also susceptible to clindamycin.

### Pathology of necrotizing enteritis

We next evaluated the intestinal histopathology of lorikeets with enteritis to determine whether we could identify *C. perfringens* in intestinal lesions. To do this, we examined formalin-fixed paraffin-embedded (FFPE) blocks of lorikeet intestinal tissue from a total of 37 CZA lorikeets that were submitted to The Ohio State University College of Veterinary Medicine between 2015 and 2019. Twenty-eight of these lorikeets had necrotizing enteritis while 7 died or were euthanized due to unrelated causes (encephalitis-2, trauma-1, non-enteric mycobacteriosis-2, air sacculitis-1, proventriculitis-1) (Table 3).

**Table 3.**
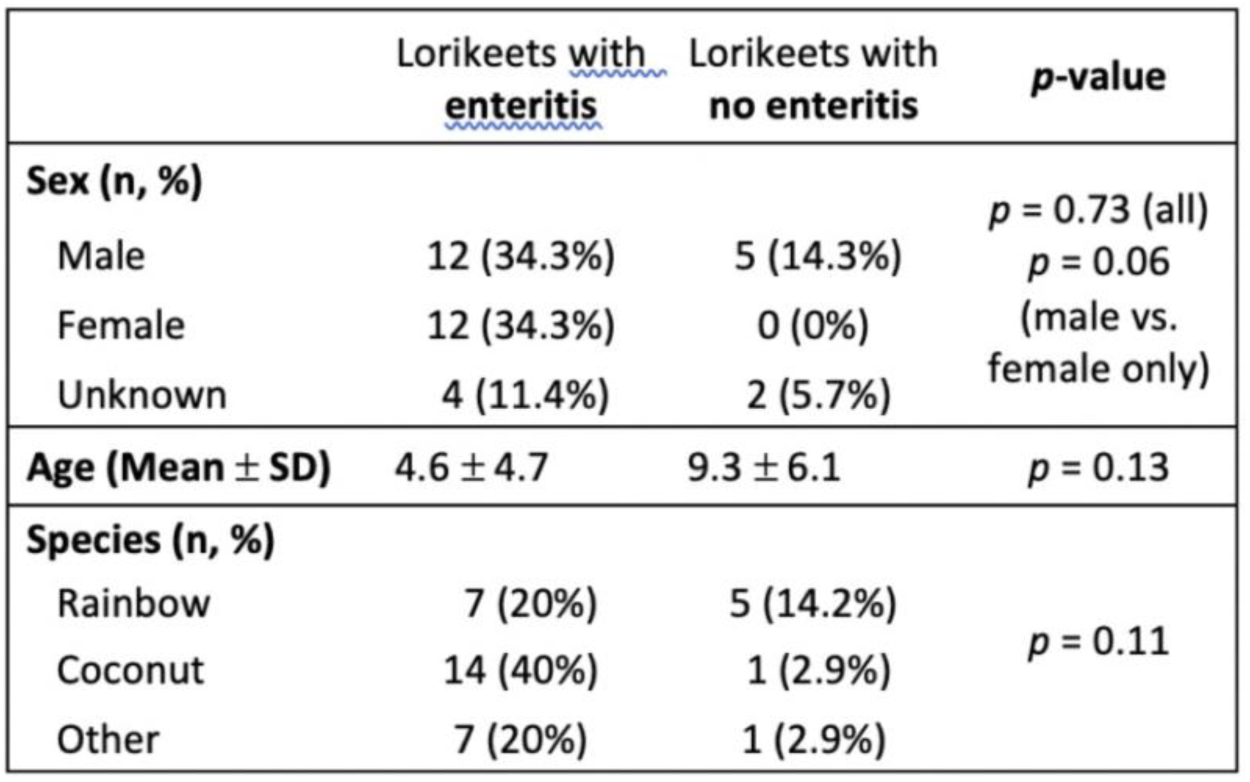
Lorikeet Demographics for CZA lorikeets submitted to pathology. The Freeman-Halton extension of the Fisher’s Exact test was used to calculate p-values for sex and species. A Kruskal-Wallis test was used to calculate the p-value for age.

There were no significant differences by sex or species in the number of lorikeets that did or did not have enteritis (by sex: Freeman-Halton extension of Fisher’s Exact Test *p =* 0.73; between males and females only, excluding “unknown”, *p* = 0.06; by species: Fisher’s Exact Test *p* = 0.11). There was also no significant difference in average age between birds with and without enteritis (Kruskal Wallis *p =* 0.13); although, birds with enteritis were generally younger. We also examined the number of enteritis deaths by month and observed an increase in cases during summer (June-September) (Fig. 3a).

**Figure 3.**
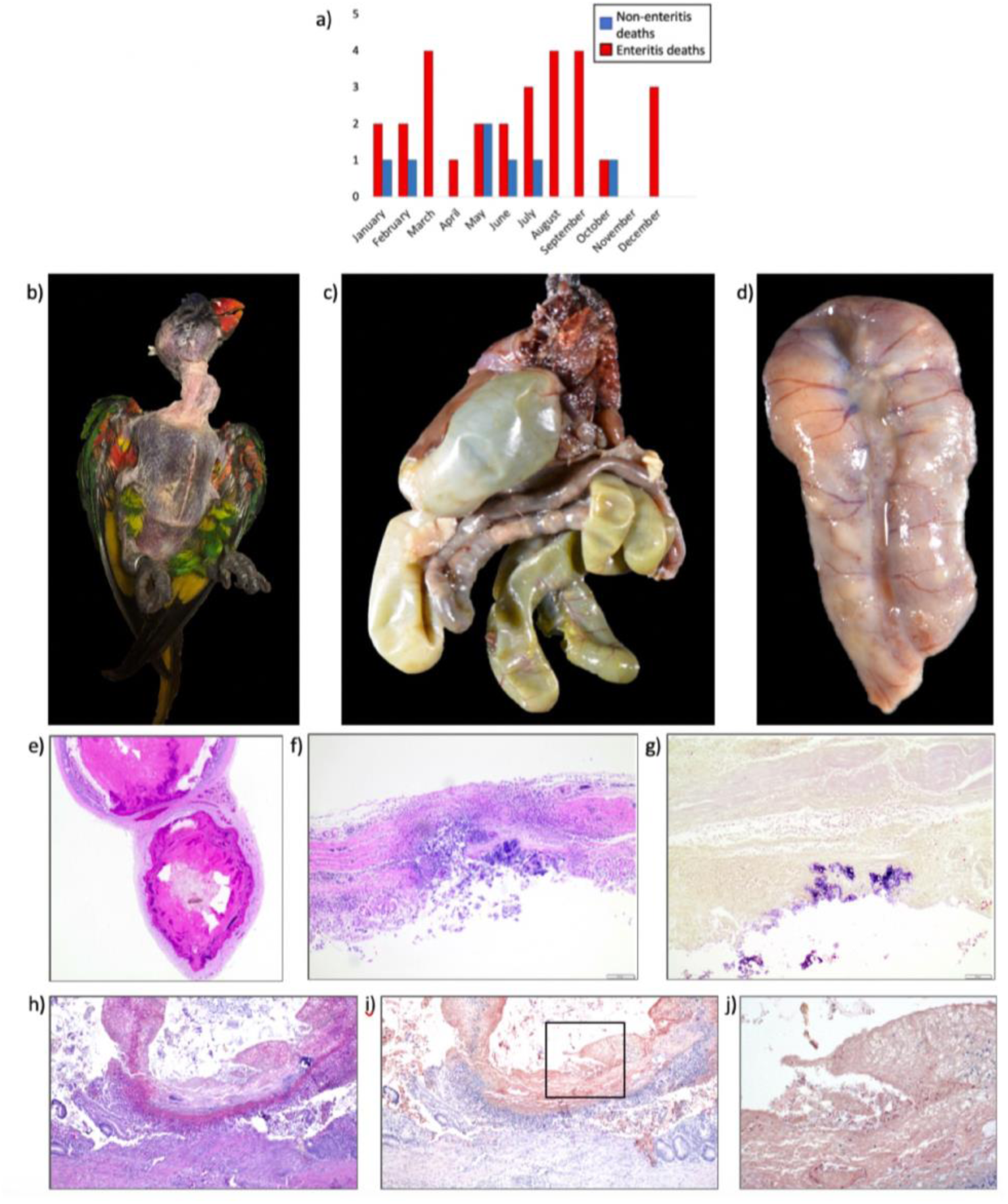
Lorikeet enteritis gross and histopathology. **a)** Number of lorikeet enteritis versus non-enteritis deaths by month. Cases from CZA were diagnosed histologically over a 5-year period (2015-2019). An increase in enteritis cases were observed during the summer months (June-September). (**b-d**) Gross necropsy findings typical of lorikeets with enteritis. **b)** The feathers have been plucked to demonstrate the decreased pectoral muscle mass of affected birds leading to a prominent keel. **c)** There are multiple severely dilated loops of intestines, including the paired ceca, with thin intestinal walls. Also present are segments of thickened and nodular intestines mottled dark tan and pale tan. Tissues are formalin fixed. **d)** Formalin fixed loop of small intestine from an affected lorikeet. There are multiple soft to firm pale tan nodules within the wall and on the serosa of the intestines. The mesentery is also thickened by pale tan tissue and similar nodules. **(e-g)** Histologic findings from lorikeets with enteritis. **e)** Hematoxylin and eosin stain (H&E) of a chronic case of necrotizing enteritis in a lorikeet. Two loops of intestine are markedly dilated with thinning of the intestinal wall and replacement with fibrosis. The lumens are impacted with a coagulum of degenerate red blood cells, bacterial colonies and sloughed mucosa that often compress the remaining atrophied and blunted intestinal villi. There is also inflammation on the serosa and adjacent mesentery. **f)** H&E of acute transmural necrotizing and ulcerative enteritis in a lorikeet. The sparse remaining mucosa is characterized by a large central ulcer, inflammatory cells including macrophages and heterophils throughout the intestinal wall centered around the ulcer and overlying large colonies of bacteria. **g)** Modified Brown-Hopps of same intestinal section as in Fig 3f. The superficially adhered bacterial colonies within the lesion are monomorphic large gram-positive bacilli. **h)** H&E of necrotizing and ulcerative enteritis in a lorikeet. There is little remaining mucosa with a large central focus of ulceration, numerous heterophils and macrophages, followed by a layer of fibrin and degenerate red blood cells with admixed large bacterial colonies and sloughed necrotic mucosal epithelium. **i)** Immunohistochemistry (IHC) against *Clostridium perfringens* from the same intestinal section as in Fig 3h. Box indicates region in Fig 3j under higher magnification. **j)** IHC against *C. perfringens* within the indicated region from Fig3i. The light brown staining is non-specific labeling of the sloughed necrotic mucosa and hemorrhage. The punctate dark brown staining indicates immunolabeling of bacteria within the necrohemorrhagic coagulum and focus of ulceration.

The majority of lorikeets had a clinical history of sudden weight loss, diarrhea, or sudden death. Most often, gross findings consisted of marked muscle wasting with a prominent keel (Fig. 3b), and multiple severely dilated loops of intestines with thin walls and scant or watery contents and/or gas (Fig. 3c). Other gross findings included tan opaque viscous coelomic fluid, intestinal segments impacted with friable dark red contents, and thickened tan mesentery with multiple small, white, firm to soft nodules within the intestines as well as on the serosa (Fig. 3d).

By histopathology, the most common finding (96 % of cases) was an intraluminal coagulum comprised of red blood cells, bacterial colonies and sloughed necrotic mucosa (Fig 3e). Approximately two thirds (64 %) of the cases had full thickness heterophilic and/or granulomatous enteritis with ulceration (Fig 3f). Approximately two thirds of the cases (61 %) had marked intestinal loop dilation, with over one third (39 %) having villous fusion and/or blunting. Thirty six percent had fibrosis within the intestinal wall, most often at sites of transmural inflammation and necrosis. The most commonly associated lesion was granulomatous and/or heterophilic coelomitis (86 %) with half of those cases having intracoelomic bacteria, most frequently bacilli or mixed bacteria. Other common lesions associated with necrotizing enteritis included mild to moderate renal tubular necrosis and/or mineralization most likely due to dehydration or septicemia (50 %) and marked extramedullary hematopoiesis within the liver (61 %). Evaluation of intestinal segments with modified Brown-Hopps Gram stain demonstrated mixed bacteria within the necrohemorrhagic coagulum and/or necroulcerative intestinal lesions. Eighty-nine percent of cases had large gram-positive bacilli present within the necrohemorrhagic coagulum and these gram-positive bacilli were also the most abundant bacteria present, followed by gram-positive cocci (71%), and gram-negative bacilli (54%) (Fig. 3g), whereas 57 % of the transmural enteritis lesions contained large gram-positive bacilli, followed by 25 % with gram-positive cocci and 21 % with gram-negative bacilli.

There was a strong correlation between the histopathologic characterization of lesions as chronic versus acute with the clinical history of repeated versus first-time enteritis cases, respectively (Fisher’s exact test, p=0.033; Relative risk 2.95, 95% CI = [0.86 – 10.08]; Sensitivity 0.875, 95% CI = [0.62 – 0.98]). The chronicity of the lesions was assessed based on the presence of fibrosis and mononuclear inflammatory infiltrates. There was no association between chronicity and the presence of *C. perfringens* (χ^2^ = 0.0009, *p =* 0.98) or *C. colinum* (χ^2^ = 0.36, *p =* 0.55).

### *Clostridium* identification and toxinotyping in FFPE lorikeet intestinal samples

We then used immunohistochemistry (IHC) and an anti-*C. perfringens* antibody to determine whether the gram-positive bacilli present in intestinal lesions were *C. perfringens*. We identified *C. perfringens* in 22 out of the 28 (79%) enteritis-positive FFPE intestinal samples and in 1 out of 7 (14%) samples that had no enteritis (Table 4, Fig. 3h,i,j). PCR Toxinotyping of the FFPE intestinal tissue identified the *C. perfringens* alpha toxin gene (*cpa*) in 13 of the 28 (46%) enteritis-positive samples and 1 of the 7 (14%) samples with no enteritis lesions. No FFPE intestinal samples were positive for any of the following toxin genes: *cpb* (beta)*, etx* (epsilon)*, itx* (iota)*, cpe* (enterotoxin), or netB (necrotic B-like). Additionally, although the gene encoding *cpb2* (beta-2 toxin) was identified in intestinal isolates, we did not find *cpb2* in the FFPE enteritis-positive samples (Table 4). *C. colinum* was identified by PCR in 18 of the 28 (84%) enteritis-positive intestines and 1 (14%) of the samples with no lesions.

**Table 4.**
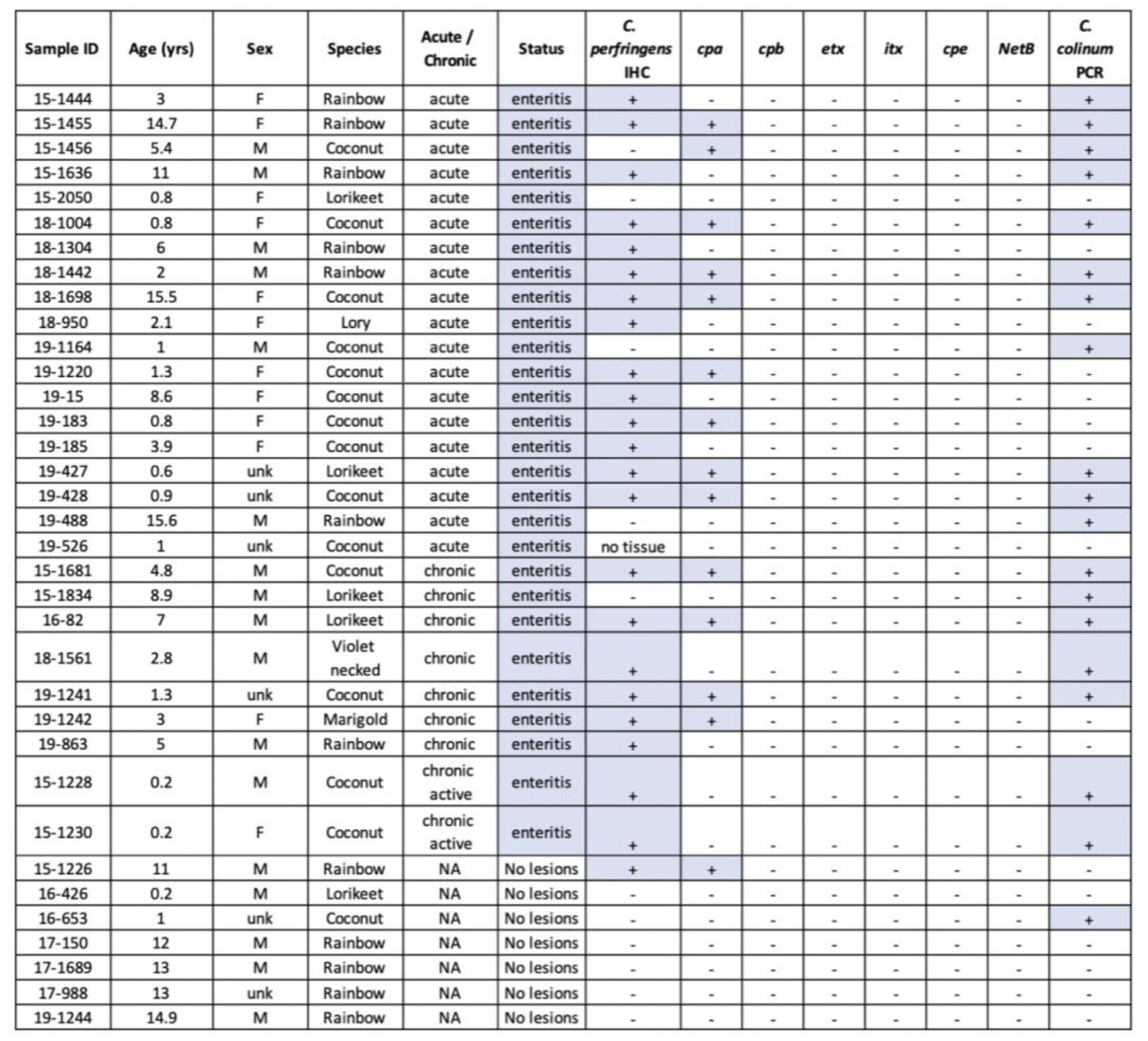
Toxin profiles, IHC, and *C. colinum* PCR on lorikeet intestines submitted to pathology.

### Denver Zoo lorikeet enteritis

To determine whether *C. perfringens* could also be found in lorikeets with enteritis at other institutions, we collected cloacal swabs and intestinal content from lorikeets at the DZ between November 2018 and May 2019. We identified 12 birds that died or were euthanized due to enteritis, and sampled these birds at necropsy (“post-mortem”). These birds were then age-, sex-, and species-matched as closely as possible to 12 healthy lorikeets that were sampled during a flock survey in May 2019 when all birds were reported to be healthy (**Additional File 10**). Similar to the CZA lorikeets, we observed decreased microbial diversity and altered microbial composition in the post-mortem lorikeets with enteritis as compared to healthy birds (Observed Features Kruskal-Wallis *p* < 0.0005, Weighted UniFrac PERMANOVA *p* = 0.001, Fig. 4**; Additional File 11**). *C. perfringens* was also significantly increased in relative abundance in the post-mortem birds (Fig. 4c,d; ANCOM W = 67). However, none of the DZ samples contained *C. colinum* 16S reads nor did they test positive for *C. colinum* via PCR. Type A *C. perfringens* was also cultured from eight of the DZ birds with enteritis.

**Figure 4.**
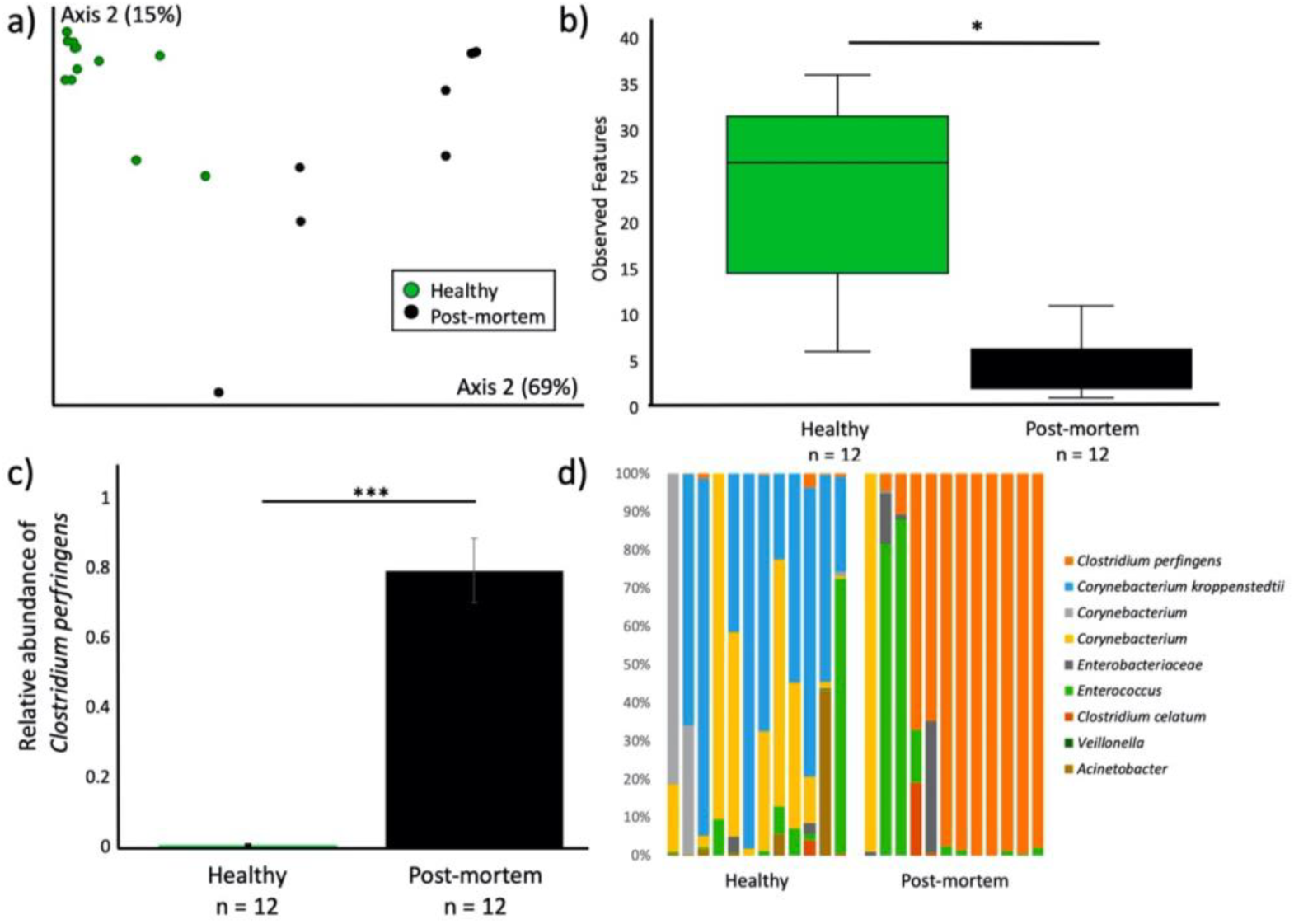
Altered microbial community diversity and composition in Denver Zoo (DZ) lorikeets with enteritis. Microbial composition and diversity in healthy lorikeets and lorikeets that died or were euthanized due to enteritis (“post-mortem” lorikeets). **a)** Microbial composition (Weighted UniFrac) was significantly altered (PERMANOVA *p* = 0.001) and **b)** microbial diversity (Observed Features) was significantly decreased (Kruskal-Wallis \**p* < 0.0005) in post-mortem lorikeets. (Also see **Additional File 11**.) **c)** The relative abundance of *C. perfringens* was also significantly increased in post-mortem lorikeets as compared to healthy lorikeets (ANCOM, W = 67). **d)** Taxa bar plots showing taxonomic distributions within healthy and post-mortem lorikeets with enteritis. Post-mortem lorikeet microbial communities were dominated by *C. perfringens*.

### Susceptibility to clostridial enteritis

#### Odds and risk ratios associated with C. perfringens and C. colinum presence in healthy birds

After identifying clostridial enteritis in two separate lorikeet flocks, we then used the CZA lorikeet data to explore factors that could be linked to susceptibility. First, we examined 16S rRNA reads for the presence of *C. perfringens* and *C. colinum* in healthy birds to determine whether they were predictors for developing future enteritis. Healthy CZA birds that had *C. perfringens* (n = 24, 36%) or *C. colinum* (n = 7, 11%) present in their gut were not at increased odds or risk of developing future enteritis (*C. perfringens*: OR = 0.59, 95% CI = [0.21-1.62], p = 0.31; RR = 0.76, 95% CI = [0.44-1.32], *p* = 0.33; *C. colinum*: OR = 1.37, 95% CI = [0.28-6.71], p = 0.69; RR = 1.16, 95% CI = [0.58-2.32], *p* = 0.67 ). Notably, *C. perfringens* and *C. colinum* were only present at low relative abundances (<1%) in healthy birds, if present at all.

#### Gut microbial composition and diversity of susceptible birds

We then divided all healthy CZA birds into two groups: True Healthy and Susceptible. True Healthy birds remained healthy throughout the entire sampling period (Nov. 2018 - Sept. 2019) and never developed enteritis. Susceptible birds were healthy birds that went on to develop *C. perfringens* enteritis at least during the sampling period. We compared the microbial communities of these groups and observed significantly increased microbial diversity and altered microbial composition in Susceptible birds as compared to True Healthy birds or birds with enteritis (Observed Features Kruskal-Wallis *p* < 0.005, Weighted UniFrac PERMANOVA *p* = 0.001, Fig. 5**; Additional File 12).**

**Figure 5.**
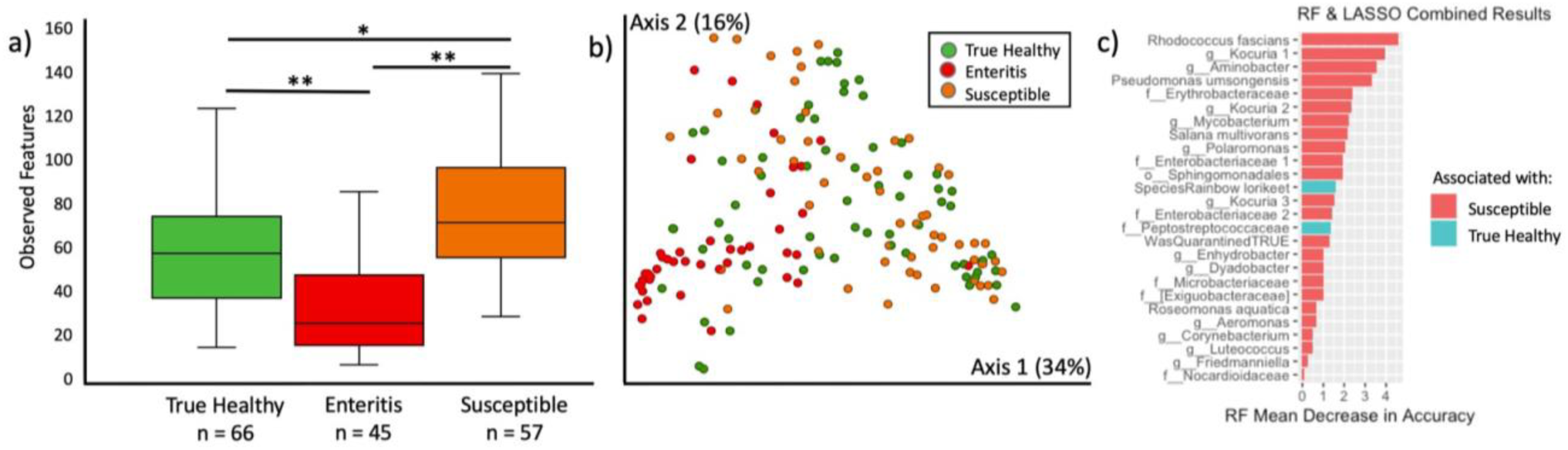
Susceptible CZA lorikeets have increased microbial diversity and altered microbial composition that predicts enteritis. Healthy lorikeets that never developed enteritis throughout the sampling period were identified as “True Healthy” while healthy birds that developed enteritis at least once during the sampling period were identified as “Susceptible.” “Enteritis” represents birds with enteritis that were sampled while they were clinically ill. No post-mortem samples are included in this figure. **a)** Microbial diversity (Observed Features, Kruskal-Wallis, all *p* <0.005) was increased and **b)** microbial composition was altered (PERMANOVA *p* =0.001) in Susceptible birds. **c)** Twenty-six variables including 24 microbial taxa and two demographic variables were identified in Random Forests and LASSO models as predictive of susceptibility or true health. The size of the bars represents the effect size of each variable as predicted by the RF model. (Also see **Additional File 12**.)

#### Predicting susceptibility using Random Forests and LASSO models

Next, we constructed Random Forests (RF) and LASSO models to compare True Healthy and Susceptible birds based on samples collected during the February 2019 CZA flock survey (all healthy birds). A total of 1503 microbial taxa (amplicon sequence variants - ASVs) were included in these models along with demographic variables including lorikeet age and species. Sex was not included as it was unknown for 31% of the birds. Seventy-five percent of the samples were used as a training set and 25% of the samples were used as a test set. The RF model identified the relative importance of variables as predictors (**Additional File 12,c**) while the LASSO model identified whether a variable was associated with susceptibility or true health (**Additional File 12,d**). We then collated variables that were identified in both the RF and LASSO models (Fig. 5c). The top 26 variables included 23 taxa associated with Susceptible birds and 1 taxon (family Peptostreptococcaceae) associated with True Healthy birds. Rainbow lorikeets (as opposed to coconut lorikeets) were also associated with health while the “WasQuarantined TRUE” variable was associated with susceptibility. This variable represented young lorikeets (< 1 year old) that were transferred from another institution; these birds underwent an initial quarantine prior to integration with the CZA flock. Some of the taxa predictive of susceptibility included: *Rhodococcus fascians*, *Kocuria* spp., *Pseudomonas umsongensis*, two taxa in the family Enterobacteriacea and an *Aeromonas* spp.

#### Dietary analysis for trypsin inhibitors

Finally, we examined lorikeet diets in relation to *C. perfringens* susceptibility. Several *C. perfringens* toxins, including *cpa*, *cpb*, *pfo*, and *cpb2* (the toxin observed in 5 CZA *C. perfringens* isolates) are sensitive to the host-produced protease trypsin [39]. However, trypsin inhibitors present in the diet can block the activity of trypsin and thereby increase the risk of *C. perfringens* toxin-mediated enteritis. As lorikeets are nectivores and their main diet under human care consists of reconstituted powdered nectar, we opted to test trypsin inhibitor levels in six commercial nectar brands including brands used at CZA and DZ. The range of trypsin inhibition for the nectars was 0-1.79 trypsin inhibitor units (TIU)/mg dry nectar, denoting relatively low inhibition (Fig. 6). For reference, raw soybeans, which have been linked to *C. perfringens* toxin-mediated enteritis in poultry, contain approximately 46 TIU/mg, and soy protein concentrate contains 9.45 TIU/mg [40]. As such, the low levels of trypsin inhibition detected in commercial nectars are unlikely to be playing a major role in susceptibility to *C. perfringens* enteritis in lorikeets; although we cannot rule out the possibility that other supplementary food items (e.g. sweet potatoes or legumes) may have contributed to toxin-mediated clostridial enteritis. Notably, *C. colinum* toxins have yet to be characterized; therefore, the role of trypsin and dietary trypsin inhibitors on *C. colinum* pathogenesis is unknown.

**Figure 6.**
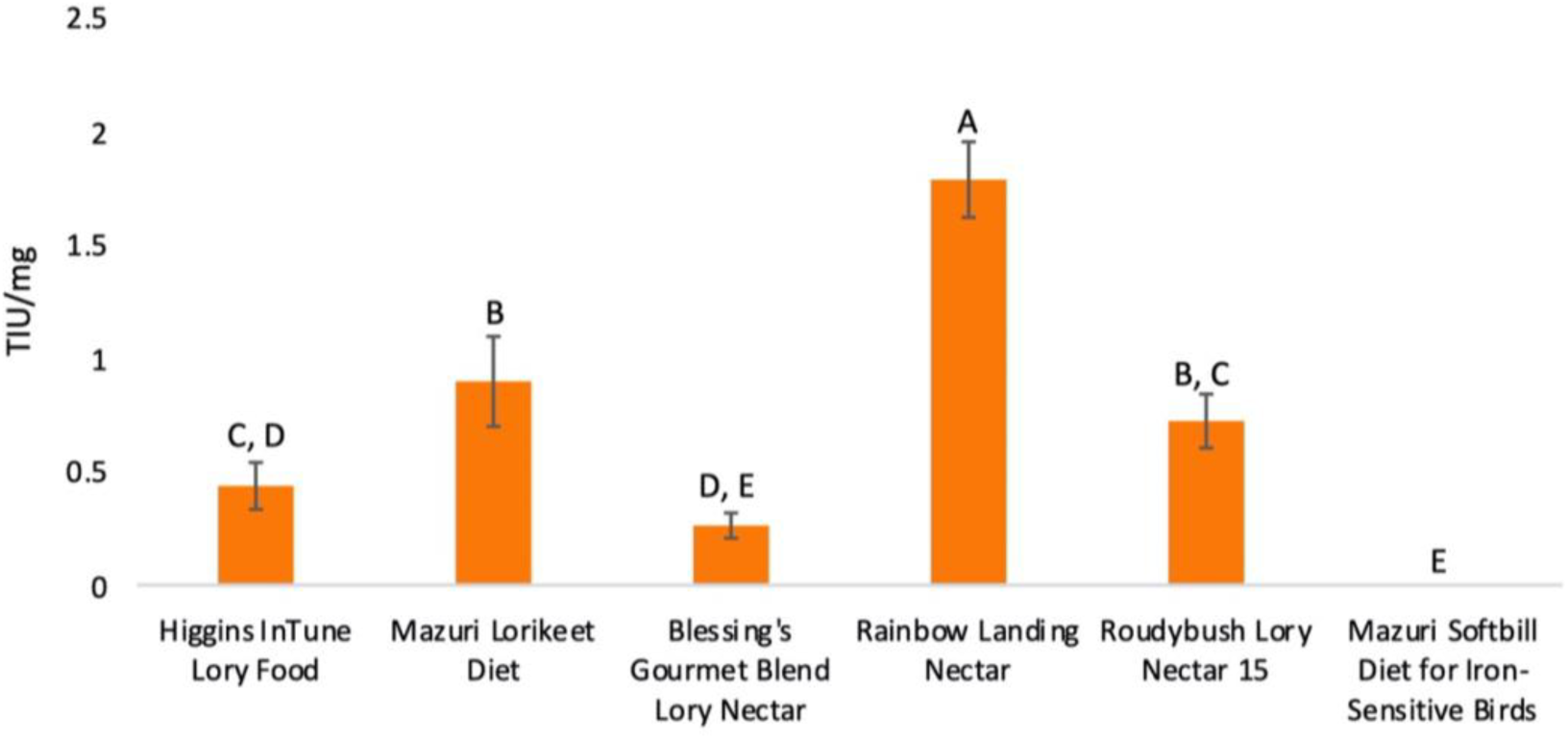
Trypsin inhibitor measurements in commercial nectars. Trypsin inhibitor units (TIU) per milligram nectar were measured in 6 commercial nectars and concentrations were compared using a one-way ANOVA and Tukey’s Test (α = 0.05). Bars that share a letter are not significantly different.

## Discussion

Our initial goal in this study was to characterize the gut microbiota of sick and healthy lorikeets with the hypothesis that a dysbiosis was driving susceptibility to enteritis. While we did identify gut microbial alterations associated with susceptibility, we also ended up identifying the probable etiologic agents of enteritis in both the CZA and the DZ lorikeet flocks. Specifically, we observed increased relative abundances of *C. perfringens* and *C. colinum* (CZA lorikeets only) in the 16S rRNA sequencing data. We then cultured lorikeet intestinal contents and identified, genotyped, and susceptibility-tested multiple *C. perfringens* isolates. A histopathologic examination of intestinal tissues further revealed inflammation, necrosis, and ulcerative lesions that also contained gram-positive bacilli consistent with clostridial enteritis and specifically *C. colinum* or *C. perfringens*. IHC and toxinotyping of intestinal tissues confirmed the presence of *C. perfringens* in lorikeets with enteritis. PCR testing also confirmed the presence of *C. colinum* in CZA lorikeets with enteritis. We then compared the gut microbiota of healthy CZA lorikeets that developed enteritis to healthy CZA lorikeets that never developed enteritis during our 10-month sampling period, and we identified several features associated with susceptibility to enteritis including: age (younger birds are more susceptible), and increased relative abundances of *Rhodococcus fascians*, *Pseudomonas umsongensis*, two taxa in the family Enterobacteriacea, and an *Aeromonas* spp., among others. This work allowed us to identify the probable causative agents of lorikeet enteritis at two zoos, develop an optimal treatment plan based on genotyping and susceptibility testing, and profile healthy birds at high risk of clostridial enteritis.

### Demographics of lorikeet enteritis

*C. perfringens* has been linked to enteritis in multiple mammal and bird species [34, 39, 41–44], including in lorikeets and other psittacines [5, 32, 33, 45–47]. In this study, young lorikeets (< 2 years old) were more likely to develop clostridial enteritis. We also observed some differences in microbial composition by age (**Additional File 5**), and age emerged as a predictor of susceptibility in the Random Forests model (**Additional File 12,c**). Previous reports in other avian species note that the immunological naivete of young birds may increase susceptibility to *C. perfringens* while adult birds are more resistant [48, 49]. We also found that coconut lorikeets were more likely to develop enteritis as compared to other lorikeet species. It is less clear what may be driving species differences in clostridial enteritis; although, there were more coconut lorikeets than any other species at CZA, and *C. perfringens* has been associated with necrotic enteritis in coconut lorikeets at other institutions [47]. While type A was the dominant *C. perfringens* toxinotype reported in the previous study on lorikeets, toxinotype C was also common. Toxinotypes B, D, E, F, and G were also observed but less common [47]. Sex has also been reported as a factor that influences susceptibility to necrotic enteritis in birds [41]; although, we did not observe differences by sex in this study. Taken together, our results suggest that both microbial and immunological factors may contribute to clostridial enteritis in young lorikeets.

### Seasonal changes in healthy lorikeet gut microbiota

We observed flock-wide shifts in gut microbiota between the three flock surveys in November 2018, February 2019, and September 2019. In previous studies, seasonal alterations in avian microbiota have been linked to diet, migration, and breeding status [21, 50–52]. Lorikeets in the CZA flock have a consistent diet and environment year-round, and they do not migrate - thus mitigating these factors as drivers for the observed seasonal changes. While breeding status could be influencing these gut microbial alterations, it is also possible that flock-wide prophylactic antimicrobial use during enteritis outbreaks and between flock surveys drove these shifts. Notably, microbial diversity decreased significantly from November 2018 to September 2019, which is a shift that could be consistent with antimicrobial use across host species including in birds [53–58].

### Lorikeet enteritis: microbes to host

Lorikeets with enteritis demonstrated microbial community shifts and histopathological changes as compared to healthy lorikeets. Based on 16S rRNA sequencing, culture, and genotyping, we confirmed the presence and increased relative abundance of *C. perfringens* type A in CZA and DZ lorikeets with enteritis. In the CZA lorikeets, we further determined that *C. perfringens* was directly associated with necroulcerative intestinal enteritis via IHC and multiple intestinal isolates of *C. perfringens* contained the *cpb2* toxin. *C. colinum* was also found at increased relative abundances in CZA but not DZ birds based on 16S sequences and PCR. In both CZA and DZ birds, we observed decreased microbial diversity and altered microbial composition in lorikeets with enteritis as compared to healthy lorikeets. *C. perfringens* and *C. colinum* have been implicated previously in necroulcerative enteritis in birds including lories, lorikeets, and poultry [47, 59–63]. Microbial community alterations have also been reported in chickens infected with *C. perfringens* [64–67]. While several clostridial species are considered normal flora in some avian species including poultry, in psittacines, clostridial species are rarely found in the intestines of healthy birds, and taxa such as *C. colinum* and *C. perfringens* are considered pathogenic [4, 31, 59, 68, 69]. Previous studies have linked the *C. perfringens* beta-2 toxin (*cpb2*) identified in CZA birds with enteritis in psittacines [33], storks [70], pigs [71], and poultry [72]; however, the cpb2 toxin has also been identified in healthy individuals (poultry, horses, dogs, and other avian species) and its role enteritis is not clear [73–77]. *C. colinum*-linked enteritis has only been reported in avian species, but virulence factors, toxins, and disease pathogenesis for *C. colinum* have yet to be fully elucidated [59].

Our results suggest that *C. perfringens* was the driver of enteritis in the DZ birds, while in the CZA birds, either *C. colinum*, or *C. perfringens*, or both acting synergistically, could have been driving the infections or creating a dysbiotic environment that allowed the other to thrive. Synergistic *C. perfringens / C. colinum* co-infections have been reported previously in poultry [78], while other clostridial co-infections (*C. perfringens* / *C. difficile*) have been reported in humans, foals, and dogs [79–81]. Phylogenetically related bacterial taxa can share functional traits and fill similar metabolic niches [82]; thus, it is feasible that an environment permissive to one type of clostridia may also be permissive to another clostridia.

### Susceptibility to enteritis

Besides age, seasonality has also been linked to necrotic enteritis in birds. In poultry, enteritis cases occur more frequently in late winter and early spring [83–85], which is what we observed in the CZA lorikeets; although historically, CZA reported enteritis cases across all seasons. DZ cases were also clustered during winter months, however this was the only occurrence of an enteritis outbreak DZ had experienced. Additionally, birds that had a history of enteritis were at 2.2 times increased risk of developing future enteritis. Gross pathology and histopathology data further linked signs of chronic lesions (fibrosis, mucosal atrophy, villus blunting/fusion, wasting) to birds with repeated cases of enteritis. Repeated bouts of enteritis in some CZA lorikeets likely resulted in permanent intestinal damage that hindered absorption and peristalsis, leading to malnutrition and intestinal stasis, which are known risk factors for *C. perfringens* enteritis [61, 86, 87]. While some lorikeets survived one or even 2 cases of enteritis, no bird lived beyond a third episode.

Diet – and specifically dietary trypsin inhibitors – can also be a risk factor for *C. perfringens* enteritis – depending on the toxinotype [45, 87, 88]. As such, we assessed trypsin inhibition in 6 commercial nectars including the nectars used at CZA and DZ. All nectars contained very low levels of trypsin inhibitors and are unlikely to be contributing to enteritis incidence. However, supplementary diet items, such as cooked sweet potatoes, which were briefly part of the DZ lorikeet diet during the enteritis outbreaks, could potentially have contained higher levels of trypsin inhibitors. Notably, trypsin inhibitor levels vary widely across sweet potato cultivars [89–91], and we did not evaluate trypsin inhibition in any supplementary foods.

Finally, we examined microbial communities in healthy birds that later went on to develop enteritis (Susceptible) or remained healthy throughout the entire study period (True Healthy). Susceptible birds displayed minor but significant differences in microbial community diversity (Observed Features), composition, and differentially abundant taxa. These microbial community differences could be linked to age as young birds were also the most susceptible. Although we found no significant difference in microbial diversity or composition in healthy lorikeets by age, birds in the youngest age group (< 2 years old) had the greatest microbial diversity (**Additional File 4, c,f**) which was also true in the Susceptible birds (Fig. 5a). Increased microbial diversity has also been observed in young chickens susceptible to *C. perfringens* infection as opposed to those that were more resistant [92]. Moreover, differences in microbial composition by age (Unweighted UniFrac, but not Weighted UniFrac) were significant in healthy birds (*p* = 0.01 **Additional File 5**). This suggests that age may influence lorikeet microbial community structure, and with a larger sample size, this may have been more apparent.

Whether shaped by age or not, the microbial community differences observed in Susceptible birds suggest that a lorikeet’s pre-existing microbial community structure could potentially influence the ability of a clostridial pathogen (e.g. *C. perfringens* or *C. colinum*) to colonize the intestinal tract. This could be achieved through alterations in the metabolic environment that create a more favorable niche for clostridia to expand. The presence of primary bile acids, for example, can act as a germinant for *Clostridium* species, while the presence of secondary bile acids (produced by bacteria that convert primary to secondary bile acids), can inhibit *C. perfringens* proliferation [93, 94]. Minor alterations in pre-existing microbial community composition and differentially abundant microbes have also been reported in chickens susceptible to *C. perfringens* [92].

We identified several microbes that were associated with susceptibility including *Rhodococcus fascians*, *Pseudomonas umsongensis*, an *Aeromonas* spp., and two taxa in the family Enterobacteriacea*. Rhodococcus fascians* has been found at increased abundances in juvenile birds (sparrows, < 1 year old) as compared to older birds and could be an age-related taxa [21]. A single human case report also highlights a co-infection between *R. fascians* and a clostridial species (*C. difficile*) [95], which leads to the intriguing question as to whether these 2 species interact in ways that may support each other’s growth. However, this co-infection was in an immunocompromised individual, so the relevance is unclear. Both *Pseudomonas* and *Aeromonas* species have been independently associated with enteritis in birds [86, 96, 97]. In a previous study that employed a subcutaneous abscess model, the addition of *Pseudomonas aeruginosa* or various Enterobacteriaceae species enhanced the growth of *C. perfringens* [98] suggesting that interactions between these taxa may indeed facilitate clostridial infections. Our RF and LASSO models also identified several other microbial taxa associated with susceptibility; although, the potential role these taxa may be playing in clostridial infections or enteritis is undetermined and requires additional study.

This study had several limitations: We identified both *C. colinum* and/or Type A *C. perfringens* in lorikeets with enteritis; however, the mechanisms by which these bacteria caused disease remain unclear. For example, while *C. colinum* has been empirically and experimentally linked to ulcerative enteritis in birds, its virulence factors have yet to be elucidated [60]. Second, although both *C. perfringens* alpha toxin (*cpa*) and beta-2 toxin (*cpb2*) have been associated with enteritis in multiple host species including birds, the role of these toxins in enteritis pathogenesis is ambiguous, and both of these toxin genes have been found in the intestines of healthy animals [39, 60, 99]. It is possible that neither *cpa* nor *cpb2* are key virulence factors in Type A. *C. perfringens* and that other unidentified virulence factors played a role in lorikeet enteritis.

Additionally, toxin gene presence (e.g. PCR, used in this study) does not necessarily equate to toxin gene expression. However, clostridia and its respective toxin genes are considered aberrant in healthy psittacines [4, 31, 59, 68, 69], suggesting that they are playing a role in enteritis even if their virulence factors are not fully defined. The surrounding gut microbiota and metabolites could also be mediating *C. perfringens* pathogenesis including colonization and toxin expression as has been demonstrated in *Clostridiodes difficile* [35, 36]

## Conclusions

In this study, we systematically examined gut microbiota and susceptibility to clostridial enteritis in two lorikeet flocks under human care. A few of our key take-aways: 1) Clostridia can be challenging to detect via culture in lorikeet cloacal swabs, but anaerobic culture of intestinal contents yielded *C. perfringens* in 6 out of 13 isolates from CZA, and 16S sequencing allowed ready identification of *C. perfringens* and *C. colinum* in birds with enteritis. As clostridia are not normal inhabitants in psittacines, this was a significant finding. 2) Clostridial isolates then underwent genotyping and susceptibility testing, which allowed us to update the lorikeets’ clinical treatment plans to more targeted therapies, aligned with antimicrobial stewardship practices (DZ – metronidazole, CZA - florfenicol and clindamycin in clinically affected birds and prophylactic flock-wide application of bacitracin). Since June of 2019, and as of this writing, there have been no new cases of enteritis in lorikeets at either CZA or DZ. 3) Young age (potentially linked with immunological naivete, limited exposures, or lower trypsin activity [100]), prior enteritis, and specific microbes including *R. fascians*, *P. umsongensis*, and Enterobacteriacea taxa are linked with susceptibility to enteritis, and these microbes could be promoting clostridial infections by establishing a niche conducive to colonization in a yet-to-be determined manner. 4) Diet – including trypsin inhibitors – can also influence susceptibility to clostridial enteritis. Although, commercial nectars were low in trypsin inhibitors, we cannot rule out the possibility that other supplementary food items (e.g. sweet potatoes or legumes) could have contributed to toxin-mediated enteritis. Clostridial enteritis, and *C. perfringens* in particular, not only affects lorikeets, but can also cause devastating losses in the poultry industry (commonly Type G *C. perfringens* with NetB toxin), and lead to gastrointestinal disease in humans and other mammals – depending on the toxinotype. This work provides a foundation for understanding gut microbiota features that are permissive to clostridial colonization and host factors (e.g. age, prior infection) that shape responses to infection.

## Methods

### Sample Collection – Columbus Zoo and Aquarium

Cloacal swabs were obtained from all healthy lorikeets at the CZA (n=67 birds) during routine flock health surveys at 3 timepoints (November 2018, February 2019, September 2019) (See **Additional File 1**, **a** for experimental design). The flock was composed of four species of lories and lorikeets including: rainbow lorikeets (*Trichoglossus moluccanus*), coconut lorikeets (*Trichoglossus haematodus*), marigold lorikeets (*Trichoglossus capistratus*), and lorys (*Trichoglossus*). Each bird was weighed and body condition scored during these surveys. Cloacal swabs, intestinal tissue, and/or intestinal contents were collected opportunistically from lorikeets (n = 34 birds) that presented with enteritis or died / were euthanized due to enteritis between November 2018 and September 2019.

A total of 246 samples were collected from birds at CZA – 172 samples from healthy birds and 74 samples from birds with enteritis. Twenty-eight additional samples were also collected from the environment and included samples of freshly prepared nectar, water, fruit, and swabs of the aviary floor and perches. Swabs from live birds that presented with enteritis were collected prior to initiation of antimicrobial therapy and, in some cases, throughout treatment. For necropsies performed at CZA, intestinal contents were milked directly into a 2 ml screw-top tube without buffer. Swabs (Puritan, Guilford, ME) and intestinal contents were immediately transferred to a - 80°C freezer and stored until sample processing.

### Sample Collection – Denver Zoo

Cloacal swabs were collected from the entire lorikeet flock during a flock survey in May 2019. At this time, all lorikeets were reported to be healthy. Cloacal swabs, intestinal tissue, and intestinal contents were collected opportunistically from lorikeets (n=12) that died or were euthanized due to enteritis between November 2018 and May 2019 (see **Additional File 1**, **b** for experimental design). These birds were then age-, sex-, and species-matched to 12 healthy lorikeets. Upon collection, all swabs and intestinal contents were immediately transferred to a - 80°C freezer and stored until sample processing and DNA extraction.

### DNA extraction, 16S rRNA amplification, and sequencing

Bacterial DNA isolation from cloacal swabs and intestinal contents (approximately 200 mg) was performed using QIAamp PowerFecal DNA Kits (Qiagen, Venlo, Netherlands). For cloacal swabs, powerbeads and lysis buffer were added directly to the screw top tubes containing the swabs. A bead beating step (6 m/s for 40 sec.) was used to replace the vortex step from the manufacturer’s protocol. The remainder of the isolation was executed according to the protocol. DNA isolation from formalin-fixed paraffin-embedded (FFPE) intestinal tissues collected at necropsy was performed using the QIAamp DNA FFPE Tissue Kit following the manufacturer recommendations. Following DNA isolation, DNA concentration was measured using a Qubit Fluorometer 4 (Invitrogen, Carlsbad, CA, USA), and purity was assessed with a NanoDrop 1000 Spectrophotometer (Thermo Scientific, Waltham, MA, USA). Samples were submitted for library preparation and sequencing at Argonne National Laboratory. Earth Microbiome Project primers 515F and 806R were used to amplify the V4 hypervariable region of the bacterial 16S rRNA gene. Amplicons were sequenced on an Illumina MiSeq in 2 x 250 paired-end mode.

### 16S rRNA sequence processing

The 16S rRNA sequences were processed, filtered, and analyzed using QIIME 2 version 2020.11 [101] and DADA2 [102]. Taxonomic assignment of amplicon sequence variants (ASVs) was performed using the Greenegenes 13_8 database with 99% sequence identity cutoff. (Note: We also performed taxonomic assignment with SILVA 132 and found that, in this case, Greengenes provided more specific taxonomic assignments, particularly in the Clostridia taxa.) A total of 246 samples from CZA and 30 samples from DZ were submitted for sequencing. Samples with fewer than 1000 reads were removed from analyses including 23 CZA samples and 6 DZ samples. This resulted in a total of 223 CZA samples and 24 DZ samples that were used in our analyses. After filtering, we obtained a total of 3,236,674 reads from the CZA samples (average: 13,911 reads per sample; range: 1003 to 68,537 reads) and 264,769 reads from the DZ samples (average: 9,026 reads per sample; range: 1922 to 26,766 reads). Sequences identified as mitochondria, chloroplasts, or eukaryotic reads were removed. Based on an examination of negative controls, we also identified the following taxa as contaminants and removed them from analyses: a taxa in the order RF39 (Mollicutes phyla); a taxa in the genus *Allobaculum*, a taxa in the genus *Massilia*; *Haemophilus parainfluenzae*; *Prevotella copri*; a taxa in the genus *Sphingomonas*; a taxa in the genus *Bradyrhizobium*; *Pseudomonas viridiflava;* and a taxa in the genus *Thermicanus*.

### Culture, bacterial identification, and genotyping

Lorikeet intestinal contents collected at necropsy were plated on reduced Trypticase Soy Agar (TSA II) with 5% sheep blood (BD BBL, Franklin Lakes, NJ) and incubated anaerobically (5% CO_2_, 5% H_2_, 90% N_2_) at 37°C until growth was seen (24 - 48 hours). As needed, bacteria was sub-cultured on *C. perfringens* selective agar, tryptose sulfite cycloserine (TSC), (Sigma Aldrich, St. Louis, MO). Bacterial colony identification was performed via Matrix Assisted Laser Desorption/Ionization Time-of-Flight Mass Spectrometry (MALDI-TOF). *C. perfringens* isolates were submitted for toxinotyping via multiplex PCR at the Ohio Department of Agriculture Animal Disease Diagnostic Laboratory (ODA ADDL). This PCR included primers for toxin types A – E (*cpa*, *cpb*, *cpb2*, *cpe*, *etx*, and *iota A*). The same primer sets and thermal cycler parameters were used to analyze DNA extracted from FFPE tissues for comparison.

### CZA Lorikeets submitted to pathology

Between 2015 - 2019, naturally deceased or humanely euthanized CZA lorikeets (due to severe clinical disease) underwent a complete macroscopic postmortem examination necropsied by veterinarians at the Columbus Zoo and Aquarium. Intestinal tracts were collected from lorikeets with enteritis (n=28) and with unremarkable intestines (n=7). Various organs, including the intestinal tracts, were placed in 10% neutral buffered formalin for fixation, stored at room temperature, and submitted to The Ohio State University College of Veterinary Medicine for evaluation of the formalin fixed organs and histopathology.

### Histopathology

Intestinal tracts, as well as other collected and formalin fixed tissues, were routinely trimmed, paraffin embedded, and stained with hematoxylin and eosin (H&E) for initial evaluation by the Comparative Pathology & Digital Imaging Shared Resource (CPDISR). Intestines were evaluated by two veterinary pathologists board-certified by the American College of Veterinary Pathologists (FU and MM) to characterize cases as necrotizing enteritis (n=28) vs control/unremarkable intestines (n=7). All sections with necrotizing enteritis were subsequently stained with the modified Brown-Hopps gram stain applied to identify and characterize intralesional bacteria, and a Masson’s Trichrome stain to characterize chronicity of the lesions through the presence of fibrosis.

### Immunohistochemistry

We then looked for the presence of *Clostridium perfringens* within intestinal sections from lorikeets with (n=28) or without enteritis (n=7) via immunohistochemistry (IHC) using a polyclonal rabbit anti-*Clostridium* perfringens antibody, OASA07164, Aviva Systems Biology, San Diego, CA). The IHC protocol is described in **Additional File 2**.

### Clostridium perfringens toxinotyping

Toxinotyping on DNA extracted from FFPE lorikeet intestinal tissues was performed at the San Bernardino branch of the California Animal Health and Food Safety (CAHFS) Laboratory, University of California-Davis, using a previously established method[47]. Additional testing for the *C. perfringens* alpha toxin (*cpa*) and beta-2 toxin (*cpb2*) was performed on CZA intestinal isolates at The Ohio State University using the following primers: cpaF (5’- GCTAATGTTACTGCCGTTGA -3’), cpaR (5’- CCTCTGATACATCG GTAAG -3’), cpb2F (5’- AGATTTTAAATATGATCCTAACC -3’) and cpb2R (5’- CAATACCCTTCACCAAATACTC -3’). PCR conditions for *cpa* and *cpb2* testing were as follows: PCR was performed in a total volume of 25 µL containing 0.75 µL of each primer (0.3 µM), 2 µL of extracted DNA, 12.5 µL of HiFi Hot Start Master Mix (Kapa Biosystems) and 9 µL of nuclease-free water. Thermocycler profiles were as follows: 95 °C for 3 min, 45 cycles of 95 °C for 30 s, 55 °C for 45 s, and 72 °C for 60 s, and a final extension step at 72 °C for 5 min. Positive controls included DNA extracted from known, toxin typed, *C. perfringens* isolates.

### Clostridium colinum PCR

PCR for *C. colinum* was also performed at CAHFS. Scrolls from (FFPE) sections of small intestine from the lorikeets were deparaffinized and the DNA extracted using a commercial kit (QIAamp DNA FFPE tissue kit; Qiagen) following the manufacturer’s instructions. The extracted DNA was used as a template for PCR amplification of a ∼192-bp fragment of the 16S rRNA gene of *C. colinum*, using the primers CcolF (5′- CGGCTGGATCACCTCCTTTC-3′) and CcolR (5′-ACATTTTTGTCTGGCTCACGA-3′). PCR was performed in a total volume of 25 µL containing 0.5 µL of each primer (0.5 µM), 3 µL of extracted DNA, 7 µL of nuclease-free water, and 14 μL of Platinum™ II Hot-Start Green PCR Master Mix (2X) (Invitrogen). Thermocycler profiles were as follows: 95 °C for 7 min, 35 cycles of 95 °C for 60 s, 60 °C for 60 s, and 72 °C for 60 s, and a final extension step at 72 °C for 7 min. Samples were held at 4 °C. Positive controls included DNA extracted from a commercial bacterial strain (ATCC 27770) and from FFPE sections of quail disease cases in which *C. colinum* had been isolated. DNA extracted from DZ samples also underwent *C. colinum* PCR.

### Measurement of trypsin inhibitor levels in nectar

We measured trypsin inhibitor levels in six commercial lorikeet feeds commonly used at zoological institutions and aviaries around North America, including at CZA and DZ. Lorikeet feeds included: Mazuri Softbill Diet for Iron-Sensitive Birds (Mazuri Exotic Animal Nutrition, St. Louis, MO), Blessing’s Gourmet Blend Lory Nectar (Blessing’s Pet Food Products, Murrieta, CA), Mazuri Lorikeet Diet (Formula: 5AB4, Mazuri Exotic Animal Nutrition, St. Louis, MO), Rainbow Landing Nectar (Berwick Productions, Inc. Escondido, CA), Roudybush Lory Nectar 15 (Roudybush, Woodland, CA), and Higgins Intune Lory Food (Higgins Premium Pet Foods, Miami, FL).

#### Nectar preparation

To eliminate assay interference due to free fatty acids, all nectars were first defatted through a hexane (Thermo Fisher Scientific, Waltham, MA) extraction. Nectars were combined with three times their volume of pure hexane and mixed for one minute. The samples were then allowed to sit for 10 minutes to allow for a separation of layers, and the top hexane-fat layer was removed. This process was repeated a total of three times for each nectar. Defatted nectars were then allowed to dry overnight in a fume hood. Once dry, 1 g of defatted nectar was added to 50 g of 0.01M NaOH (Thermo Fisher Scientific, Waltham, MA). The mixture was stirred slowly on a stir plate for 3 hours. Extracts were then centrifuged at 4696 x g, and the supernatant was decanted to produce the final extract.

#### Trypsin inhibitor assay

Trypsin inhibitor assays were carried out based on standard American Association of Cereal Chemists (AACC) methods [103], with modifications as proposed by Liu (2019) [40]. Reagent preparation is described in **Additional File 3**. Nectar solutions were diluted to various levels by combining 0 - 1 mL of nectar extract with enough deionized water to yield 2 mL total. Concentration ranges chosen were based upon preliminary trials, with the aim of yielding absorbance data points that fell in the range of 30 – 70 % inhibition. The 2 mL of diluted nectar were added to 15-mL centrifuge tubes and combined with 5 mL benzoyl-DL-arginine-p-nitroanilide hydrochloride (BAPA) solution. The mixture was incubated in a 37 °C water bath to bring it up to temperature, and then the assay reaction was initiated by adding 2 mL pre-warmed trypsin solution (0.02 mg trypsin/mL) to each tube. The tubes were allowed to react for exactly 10 minutes at 37 °C, then the reaction was stopped with the addition of 1 mL 30 % acetic acid solution. Samples were allowed to cool to room temperature before measuring absorbance at 410 nm with an HP 8453 UV-Vis spectrophotometer (Hewlett Packard, Palo Alto, CA). Absorbance readings were corrected with nectar blanks by mixing all reagents, but adding trypsin solution after the acetic acid to ensure the enzyme was inactive. A positive control sample was also made using 2 mL water in place of nectar and running the assay as delineated above.

#### TIU calculation

With the definition that 1 TIU = a decrease in 0.01 absorbance compared to a positive control sample, TIU/mg could be calculated as follows:

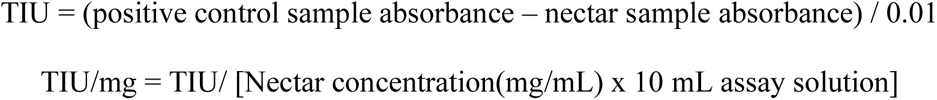

To compare trypsin inhibitor concentrations (TIU/mg) between nectars, we applied a one-way ANOVA followed by pairwise Tukey’s tests.

### Statistical analyses

We compared the number of lorikeets that ever had enteritis versus the number of lorikeets that never developed enteritis by age, sex, and species using a χ^2^ test [104]. In cases where groups had a frequency less than 5, we use the Yates’ χ^2^ correction. For cases in which a group contained zero individuals (e.g. 0 females), we used the Freeman-Halton extension of the Fisher’s exact test. To compare average age across groups, we used a Kruskal-Wallis test after testing for normality using a Shapiro-Wilk test. For microbial community analyses, alpha diversity was compared between groups using observed features, the Shannon diversity metric, and the Kruskal-Wallis test. Beta diversity was evaluated using permutational multivariate analysis of variances (PERMANOVAs) between groups on Bray-Curtis distance matrices. All alpha and beta diversity *p*-values were corrected for multiple comparisons using the Benjamini Hochberg false discovery rate (FDR) correction. A *p-*value < 0.05 was considered significant. Differential abundances of microbes by status (healthy, enteritis, susceptible) or season (November 2018, February 2019, September 2019) were tested using an ANCOM [105].

Machine learning methods were used to assess susceptibility to enteritis based on microbial relative abundances and other demographic factors. Specifically, microbial composition was used in a supervised setting for classifying birds into True Healthy and Susceptible groups. True Healthy birds never developed enteritis throughout the sampling period (Nov. 2018 - Sept. 2019) while Susceptible birds developed enteritis at least once during this time. Random forests (RF)[106] and logistic regression with appropriate regularization (LASSO)[107, 108] were employed to differentiate these groups. The predictive accuracy was then assessed through cross-validation using an area under the receiver-operating characteristics curve (ROC).

## Declarations

### Ethics approval and consent to participate

IACUC approval through The Ohio State University (# 2019A00000028).

### Consent for publication

Not applicable.

### Availability of data and material

Sequencing data is available at NCBI Bioproject PRJNA722436.

### Funding

Funding for this project provided by the following:

Columbus Zoo/Ohio State Cooperative Grants Program

National Institutes of Health Training Grant (T35)

The Ohio State University Infectious Diseases Institute

The Ohio State University College of Veterinary Medicine

American Association of Zoo Veterinarians’ Wild Animal Health Fund

The Comparative Pathology & Digital Imaging Shared Resource is supported in part by grant P30 CA016058 from the National Cancer Institute, Bethesda, MD.

### Authors’ contributions

RJ and JH were involved in identification of the clinical syndrome, project conceptualization, sample collection, and clinical data.

DM, CM, and VH were involved in project development, DNA extraction, sequencing data analysis and interpretation, manuscript writing, and figure preparation.

VH also obtained project support and oversaw/managed the project and performed demographic / epidemiological analyses. RMrochak, RMadan, KS, and CM were involved in toxinotyping on intestinal isolates. ME was involved in sequencing data filtering and processing.

LG and SP were involved in Random Forests and LASSO analysis and interpretation, manuscript writing, and figure preparation. KF-K, MM, KLP, GB, and FU were involved in histopathology analysis, staining, and interpretation. MM and FU were also involved in manuscript writing and figure preparation. KF-K and MM were involved in gross pathology interpretation and figure preparation. FU performed histopathology, IHC analysis, interpretation, and figure preparation. MN performed toxinotyping on FFPE blocks, *C. colinum* PCR, and was involved in manuscript writing and table preparation. WC, KS, and YV were involved in trypsin inhibitor testing and data interpretation. WC and YV were also involved manuscript writing and figure preparation.

## Acknowledgements

We would like to acknowledge all of the keepers and veterinary staff at the Columbus Zoo and Aquarium and the Denver Zoo for facilitating sample collection and providing detailed health, medication, and dietary record. We also thank Dr. Michael Martinez for assistance with gross pathology photos, and we gratefully acknowledge Drs. Tamara Kruse and Helena Mendes-Soares for early discussions that shaped this project.

## Additional Files

**Additional File 1: Experimental design.** Lorikeet sampling by season and opportunistically during cases of enteritis at the a) Columbus Zoo and Aquarium and the b) Denver Zoo.

**Additional File 2: *Clostridium perfringens* IHC protocol from California Animal Health & Food Safety Laboratory**

**Additional File 3: Reagent preparation for measurement of Trypsin Inhibitor Levels in nectar**

**Additional File 4: Microbial community diversity in healthy CZA birds by sex, species, and age.** Healthy birds across all time points: Microbial diversity (Shannon Diversity Index and Observed Features) did not differ significantly by **a**,**d**) sex, **b**,**e**) species, or **c**,**f**) age (Kruskal Wallis: Shannon sex *p* = 0.77, species *p* = 0.25, age *p* = 0.89; Observed Features sex *p* = 0.93, species *p* = 0.18, age *p* = 0.08).

**Additional File 5: CZA Microbial community composition by sex, species, and age.**

**Additional File 6: Seasonality in gut microbiota in healthy CZA lorikeets. a)** Microbial community composition (Unweighted UniFrac) and **b)** diversity (Shannon Diversity Index) by season (November 2018, February 2019, September 2019). There were significant shifts in microbial composition by season (PERMANOVA *p* = 0.001), and microbial diversity decreased significantly between November 2018 and September 2019 (Kruskal-Wallis \**p* < 0.001). (Also see Fig. 1.)

**Additional File 7: Altered microbial diversity and composition in CZA lorikeets with enteritis.** Microbial composition and diversity in healthy lorikeets, lorikeets with enteritis, and lorikeets that died or were euthanized due to enteritis (post-mortem). **a)** Microbial composition (Unweighted UniFrac) was significantly altered (PERMANOVA *p* = 0.001) and **b)** microbial diversity (Shannon Diversity Index) was significantly decreased (Kruskal-Wallis \**p* < 0.005, \*\**p* < 0.0005, \*\*\**p* < 0.00001) in lorikeets with enteritis or post-mortem lorikeets. (Also see Fig. 2.)

**Additional File 8: Differentially abundant microbes by health status.** Based on an ANCOM, Clostridia were significantly increased in relative abundance in lorikeets with enteritis as compared to healthy lorikeets. This analysis included 157 healthy samples and 45 enteritis samples from a total of 67 birds. No post-mortem samples were included in this analysis. Clostridia are in **bold** text.

**Additional File 9: Differentially abundant microbes by health status in 25 birds with matched healthy / enteritis samples.** Based on an ANCOM, Clostridia were significantly increased in relative abundance in lorikeets with enteritis as compared to healthy lorikeets. This analysis included 29 healthy samples and 35 enteritis samples from a total of 25 birds. No post-mortem samples were included in this analysis. Clostridia are in **bold** text.

**Additional File 10: Denver Zoo (DZ) Lorikeet Demographics.** The p-value for sex were based on a χ^2^ test. A Kruskal-Wallis test was used to calculate the p-value for age.

The Freeman-Halton extension of the Fisher’s Exact test was used to calculate a p-value for species.

**Additional File 11: Altered microbial composition and diversity in Denver Zoo lorikeets with enteritis.** Microbial composition and diversity in healthy lorikeets and lorikeets that died or were euthanized due to enteritis (post-mortem). **a)** Microbial composition (Unweighted UniFrac) was significantly altered (PERMANOVA *p* = 0.001) and **b)** microbial diversity was significantly decreased (Shannon, Kruskal-Wallis \**p* < 0.005) in post-mortem lorikeets. (Also see Fig. 4.)

**Additional File 12: Susceptible CZA lorikeets have altered microbial composition that predicts enteritis.** Healthy lorikeets that never developed enteritis throughout the sampling period were identified as “True Healthy” while healthy birds that developed enteritis at least once during the sampling period were identified as “Susceptible.” “Enteritis” represents birds with enteritis that were sampled while they were clinically ill. No post-mortem samples are included in this figure. **a)** Microbial diversity (Shannon, Kruskal-Wallis, *p* < 0.005) was increased in Susceptible and True Healthy birds as compared to birds with enteritis and **b)** microbial composition (Unweighted UniFrac) was altered in Susceptible birds. Variables associated with susceptibility or health were then predicted by a **c)** Random Forest (RF) or **d)** LASSO model. The RF model has a sensitivity of 0.75, a specificity of 0.571, and an overall accuracy of 0.557. This model identifies the relative importance of each variable but not whether the variable is associated with susceptibility or health. The LASSO model has a sensitivity of 0.875, a specificity of 0.571, an overall accuracy of 0.733, and generates an area under the curve (AUC) of 0.72. This model (LASSO) identifies whether a variable is associated with susceptibility or health but not the relative importance of the variable.

