## Additional Files for "Gut microbiota and age shape susceptibility to clostridial enteritis in lorikeets under human care"

**Additional File 1: Experimental design.** Lorikeet sampling by season and opportunistically during cases of enteritis at the a) Columbus Zoo and Aquarium and the b) Denver Zoo.

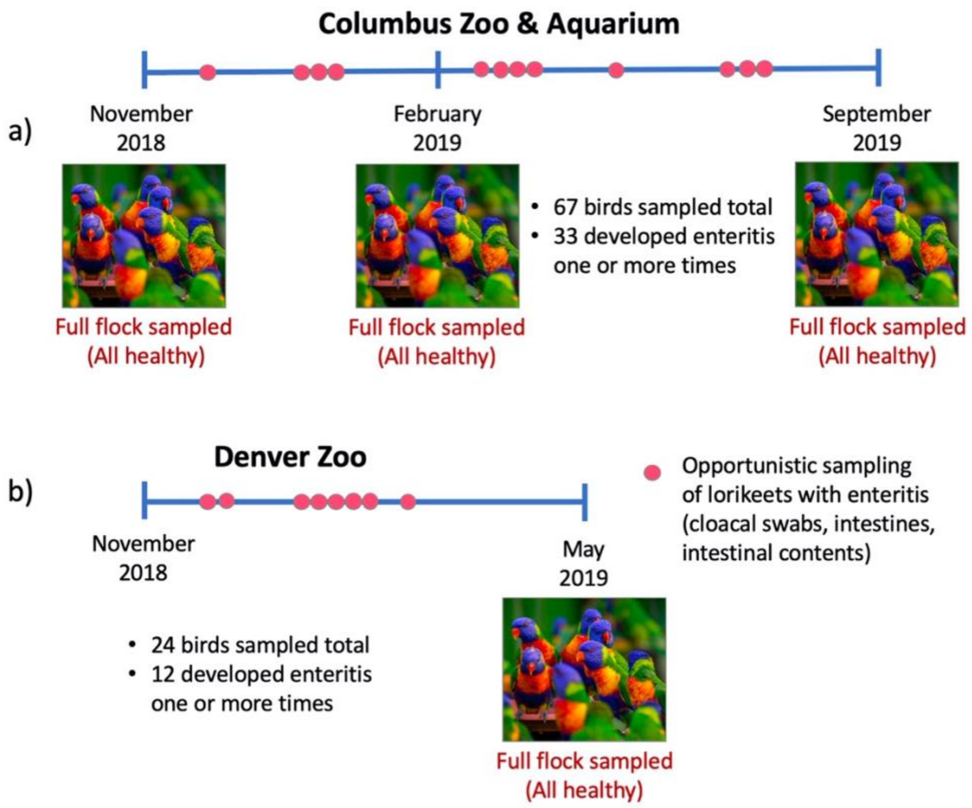

**Additional File 2: *Clostridium perfringens* IHC protocol from California Animal Health & Food Safety Laboratory**

| | CALIFORNIA ANIMAL HEALTH & FOOD SAFETY LABORATORY | | Page 1 of 3<br>Revision #: 4<br>Status: Active |
|---|---|---|---|
| **Document #:** | SHIS-02-123 | **Effective date:** | *June 13, 2019* |
| **Document Title:** | Clostridium perfringens testing in paraffin-embedded tissue sections | **Supersedes:** | SHIS-02-123-3 |
| **Author:** | Juliann Saputo | | |

Controlled hard copies of this document are marked on the first page with a colored stamp

**PURPOSE**

To detect Clostridium perfringens in paraffin-embedded tissue sections.

**SCOPE**

Clostridium perfringens antibodies are labeled with a peroxidase labeled polymer and visualized with a colored substrate-product reaction.

**DEFINITIONS AND ACRONYMS**

1. IHC: Immunohistochemistry
2. TBS: Tris buffered saline

**SPECIMEN INFORMATION**

Formalin fixed, paraffin-embedded tissues.

**REAGENTS AND MEDIA**

1. Antigen Retrieval Solution
   Distilled Water ----------------------------------------------------150 ml
   Pepsin ------------------------------------------------------------ .6 gm
   Hydrochloric acid ----------------------------------------------- 1.5 ml
2. Endogenous Peroxidase Quenching Solution, 3%
   Distilled Water ---------------------------------------------------135 ml
   Hydrogen peroxide (30%) ------------------------------------- 15 ml
3. Rinse Buffer pH 7.4
   TBS Autowash Buffer-20x(or equivalent)-------------------50ml
   Commercial Product-BioCare Medical
   Distilled Water ---------------------------------------------------- 950 ml
4. Antibody Diluent
   DaVinci Green Diluent-or equivalent---------------------RTU
5. Blocking Solution
   Background Punisher or equivalent ---------------------------RTU
   Commercial Product-BioCare Medical
6. Primary Antibody Solution
7. Secondary Detection Solution
   Anti-rabbit polymer labeled with HRP or equivalent--------RTU
8. Nova Red AEC Substrate Chromagen Kit or equivalent---Kit
9. Mayer's Hematoxylin (commercial)

All reagents are applied at 200-250ul per slide.

Uncontrolled

### Additional File 3: Reagent preparation for measurement of Trypsin Inhibitor Levels in nectar

#### Methods

##### *Reagent preparation*

A stock solution of Tris buffer was made by adding 6.05 g Tris (Sigma-Aldrich, St. Louis, MO) with 2.94 g calcium chloride ($CaCl_2$) (Sigma-Aldrich, St. Louis, MO) to approximately 20 mL deionized water. The pH of the solution was adjusted to 8.2 with 1N hydrochloric acid (Thermo Fisher Scientific, Waltham, MA), then the solution was brought up to a total volume of 50 mL with water. To make the working solution, 5 mL Tris stock solution was combined with deionized water to a final volume of 100 mL (1:20 dilution). A trypsin stock solution was produced by adding 20 mg bovine trypsin (Sigma-Aldrich, St. Louis, MO) to 50 mL 0.001M HCl. To make the working solution, a 1:20 dilution was performed to yield a concentration of 0.02 mg trypsin/mL. The substrate solution was prepared by mixing 40 mg benzoyl-DL-arginine-p-nitroanilide hydrochloride (BAPA) (Sigma-Aldrich, St. Louis, MO) with 1 mL dimethyl sulfoxide (Sigma-Aldrich, St. Louis, MO). The mixture was gently heated on a hot plate at 32◦C to aid with dissolution before being added to 100 mL Tris buffer. This solution was incubated in a Precision SWB 15, Model TSSWB15 water bath (Thermo Scientific, Waltham, MA) set at 37◦C. A 30% acetic acid solution was made by combining 30 mL glacial acetic acid (Thermo Fisher Scientific, Waltham, MA) with 70 mL deionized water.

**Additional File 4: Microbial community diversity in healthy CZA birds by sex, species, and age.** Healthy birds across all time points: Microbial diversity (Shannon Diversity Index and Observed Features) did not differ significantly by **a**,**d**) sex, **b**,**e**) species, or **c**,**f**) age (Kruskal Wallis: Shannon sex $p = 0.77$, species $p = 0.25$, age $p = 0.89$; Observed Features sex $p = 0.93$, species $p = 0.18$, age $p = 0.08$).

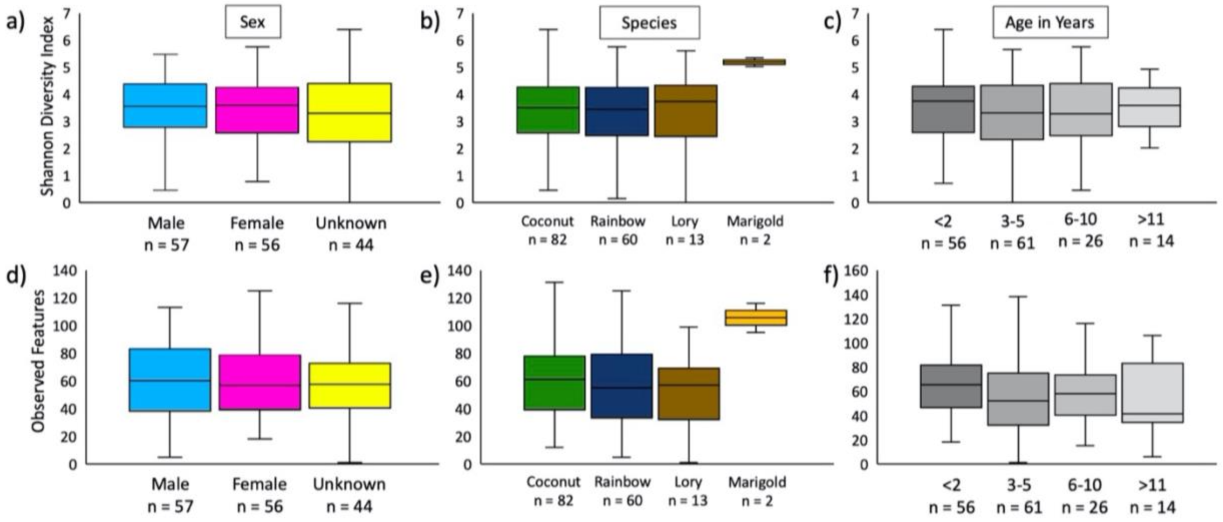

### Additional File 5: CZA Microbial community composition by sex, species, and age.

**Additional File 5: CZA microbial community composition by sex, species, and age.**

| Beta-diversity metric | Category | PERMANOVA *p*-value |
|---|---|---|
| Weighted UniFrac | Sex | 0.26 |
| Unweighted UniFrac | Sex | 0.05 |
| Weighted UniFrac | Species | 0.32 |
| Unweighted UniFrac | Species | 0.35 |
| Weighted UniFrac | Age | 0.47 |
| Unweighted UniFrac | Age | 0.01 |

**Additional File 6: Seasonality in gut microbiota in healthy CZA lorikeets. a)** Microbial community composition (Unweighted UniFrac) and **b)** diversity (Shannon Diversity Index) by season (November 2018, February 2019, September 2019). There were significant shifts in microbial composition by season (PERMANOVA $p = 0.001$), and microbial diversity decreased significantly between November 2018 and September 2019 (Kruskal-Wallis *$p < 0.001$). (Also see **Fig. 1**.)

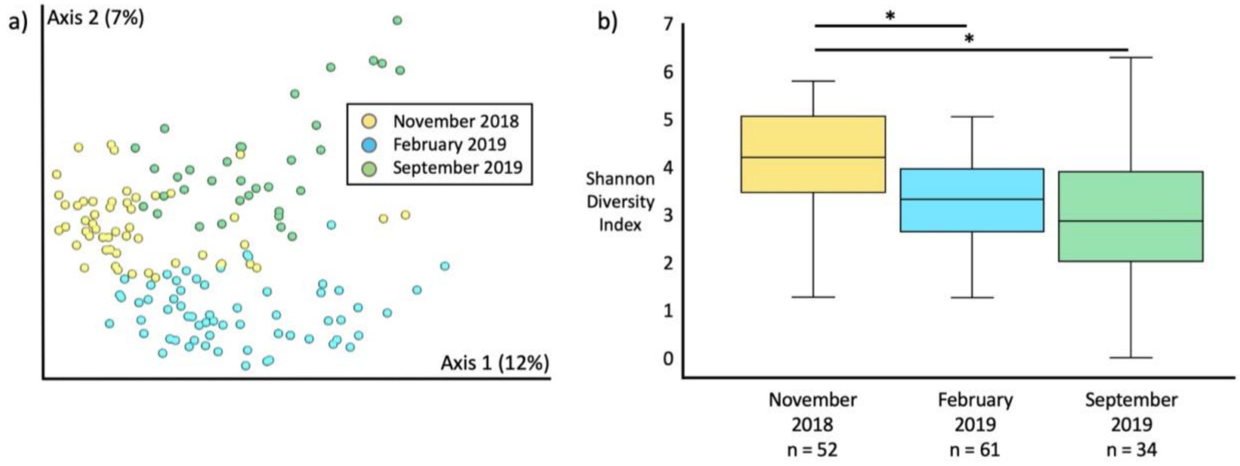

**Additional File 7: Altered microbial diversity and composition in CZA lorikeets with enteritis.** Microbial composition and diversity in healthy lorikeets, lorikeets with enteritis, and lorikeets that died or were euthanized due to enteritis (post-mortem). **a)** Microbial composition (Unweighted UniFrac) was significantly altered (PERMANOVA $p = 0.001$) and **b)** microbial diversity (Shannon Diversity Index) was significantly decreased (Kruskal-Wallis *$p < 0.005$, **$p < 0.0005$, ***$p < 0.00001$) in lorikeets with enteritis or post-mortem lorikeets. (Also see **Fig. 2.**)

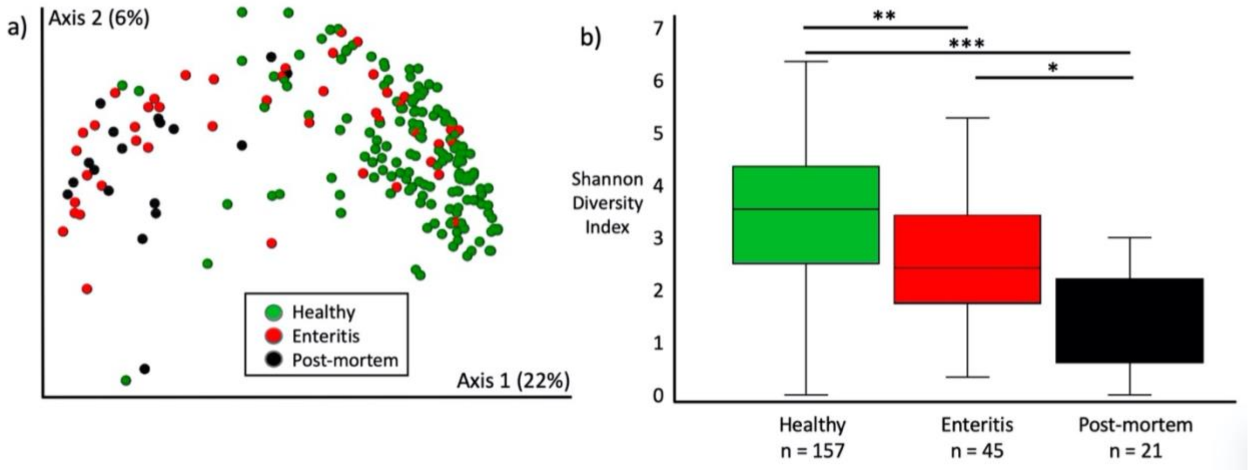

**Additional File 8: Differentially abundant microbes by health status.** Based on an ANCOM, Clostridia were significantly increased in relative abundance in lorikeets with enteritis as compared to healthy lorikeets. This analysis included 157 healthy samples and 45 enteritis samples from a total of 67 birds. No post-mortem samples were included in this analysis. Clostridia are in **bold** text.

**Additional File 8: Differentially abundant microbes by health status.** Based on an ANCOM, Clostridia were significantly increased in relative abundance in lorikeets with enteritis as compared to healthy lorikeets. This analysis included 157 healthy samples and 45 enteritis samples from a total of 67 birds. No post-mortem samples were included in this analysis. Clostridia are in **bold** text.

| ASV | Taxa | W value | Increased in Enteritis |
|---|---|---|---|
| a3000823e9ab005bb353ff4e1e20eed8 | **p__Firmicutes; c__Clostridia; o__Clostridiales; f__Clostridiaceae; g__Clostridium; s__perfringens** | 1098 | **Yes** |
| 89dc0dab78a4513dac4e8f45c5bb1cfa | **p__Firmicutes; c__Clostridia; o__Clostridiales; f__Clostridiaceae; g__Clostridium; s__neonatale** | 1094 | **Yes** |
| bd4017ad4efac59720e2d164da18ace4 | **p__Firmicutes; c__Clostridia; o__Clostridiales; f__Clostridiaceae; g__Clostridium; s__paraputrificum** | 1094 | **Yes** |
| 91db31e427ef7bc5e5b8480e19ee397f | p__Firmicutes; c__Bacilli; o__Lactobacillales; f__Lactobacillaceae; g__Lactobacillus; s__salivarius | 1094 | Yes |
| b53845c93c742c75c18148e47303150d | p__Firmicutes; c__Clostridia; o__Clostridiales; f__Lachnospiraceae | 1082 | Yes |
| 87109f2ecbac128515490ec559d6fa2c | p__Firmicutes; c__Bacilli; o__Bacillales; f__Staphylococcaceae; g__Staphylococcus | 1081 | Yes |
| 8a31169dff6a8c2d1e5f36f68025bd74 | **p__Firmicutes; c__Clostridia; o__Clostridiales; f__Lachnospiraceae; g__Clostridium; s__colinum** | 1077 | **Yes** |
| fc4f9797735349ed7612b11e73a1b4da | p__Firmicutes; c__Bacilli; o__Lactobacillales; f__Lactobacillaceae; g__Lactobacillus; s__salivarius | 1077 | Yes |
| 96146193779c38c68a78bdb867afc0b1 | **p__Firmicutes; c__Clostridia; o__Clostridiales; f__Lachnospiraceae; g__Clostridium; s__colinum** | 1043 | **Yes** |
| cd42629a00d89a92cd2e52733d7b3682 | **p__Firmicutes; c__Clostridia; o__Clostridiales; f__Clostridiaceae; g__Clostridium; s__neonatale** | 1043 | Yes |
| 28c4ab46bcdb4e950e0ef2b6c7142cf2 | p__Proteobacteria; c__Gammaproteobacteria; o__Enterobacteriales; f__Enterobacteriaceae | 1043 | No |
| 66a18cad718309ba7c8ee1199b12eb85 | p__Actinobacteria; c__Actinobacteria; o__Actinomycetales; f__Micrococcaceae; g__Kocuria; s__ | 1021 | No |
| 1878459013cf15f2993a81c14978c980 | p__Firmicutes; c__Bacilli; o__Bacillales; f__Staphylococcaceae; g__Staphylococcus | 1017 | No |
| 119c7765967bf29001a0b869513c5879 | p__Actinobacteria; c__Actinobacteria; o__Actinomycetales; f__Micrococcaceae; g__Kocuria | 1014 | No |

**Additional File 9: Differentially abundant microbes by health status in 25 birds with matched healthy / enteritis samples.** Based on an ANCOM, Clostridia were significantly increased in relative abundance in lorikeets with enteritis as compared to healthy lorikeets. This analysis included 29 healthy samples and 35 enteritis samples from a total of 25 birds. No post-mortem samples were included in this analysis. Clostridia are in **bold** text.

**Additional File 9: Differentially abundant microbes by health status in 25 birds with matched healthy / enteritis samples.** Based on an ANCOM, Clostridia were significantly increased in relative abundance in lorikeets with enteritis as compared to healthy lorikeets. This analysis included 29 healthy samples and 35 enteritis samples from a total of 25 birds. No post-mortem samples were included in this analysis. Clostridia are in **bold** text.

| ASV | Taxa | W value | Increased in Enteritis |
|---|---|---|---|
| a3000823e9ab005bb353ff4e1e20eed8 | **p__Firmicutes; c__Clostridia; o__Clostridiales; f__Clostridiaceae; g__Clostridium; s__perfringens** | 788 | **Yes** |
| 89dc0dab78a4513dac4e8f45c5bb1cfa | **p__Firmicutes; c__Clostridia; o__Clostridiales; f__Clostridiaceae; g__Clostridium; s__neonatale** | 779 | **Yes** |
| e4e259a27d64e0d63fc88ebcc68de3e5 | p__Actinobacteria; c__Actinobacteria; o__Actinomycetales; f__Corynebacteriaceae; g__Corynebacterium; s__ | 752 | No |
| bcb4afaffb8f31c47c067f20e9ad9a34 | p__Proteobacteria; c__Gammaproteobacteria; o__Pseudomonadales; f__Pseudomonadaceae; g__Pseudomonas; s__veronii | 747 | No |
| 119c7765967bf29001a0b869513c5879 | p__Actinobacteria; c__Actinobacteria; o__Actinomycetales; f__Micrococcaceae; g__Kocuria | 735 | No |
| 28c4ab46bcdb4e950e0ef2b6c7142cf2 | p__Proteobacteria; c__Gammaproteobacteria; o__Enterobacteriales; f__Enterobacteriaceae | 728 | No |
| 5fcc3fa2e888c52fe2bc176365ad60ad | p__Proteobacteria; c__Gammaproteobacteria; o__Enterobacteriales; f__Enterobacteriaceae | 718 | No |

**Additional File 10: Denver Zoo (DZ) Lorikeet Demographics.** The p-value for sex were based on a $\chi^2$ test. A Kruskal-Wallis test was used to calculate the p-value for age.

The Freeman-Halton extension of the Fisher's Exact test was used to calculate a p-value for species.

**Additional File 10: Denver Zoo (DZ) Lorikeet Demographics.** The p-value for sex were based on a $\chi^2$ test. A Kruskal-Wallis test was used to calculate the p-value for age. The Freeman-Halton extension of the Fisher's Exact test was used to calculate a p-value for species.

| | Lorikeets with **enteritis** | Lorikeets with **no enteritis** | ***p*-value** |
|---|---|---|---|
| **Sex (n, %)** | | | $p$ = 0.41 |
| Male | 5 (21%) | 7 (29%) | |
| Female | 7 (29%) | 5 (21%) | |
| **Age (Mean ± SD)** | 6.3 ± 4.3 | 5.9 ± 4.1 | $p$ = 0.89 |
| **Species (n, %)** | | | $p$ = 0.67 |
| Rainbow | 7 (29%) | 9 (37.5%) | |
| Coconut | 4 (17%) | 3 (12.5%) | |
| Other | 1 (4%) | 0 (0%) | |

**Additional File 11: Altered microbial composition and diversity in Denver Zoo lorikeets with enteritis.** Microbial composition and diversity in healthy lorikeets and lorikeets that died or were euthanized due to enteritis (post-mortem). **a)** Microbial composition (Unweighted UniFrac) was significantly altered (PERMANOVA $p = 0.001$) and **b)** microbial diversity was significantly decreased (Shannon, Kruskal-Wallis *$p < 0.005$) in post-mortem lorikeets. (Also see **Fig. 4**.)

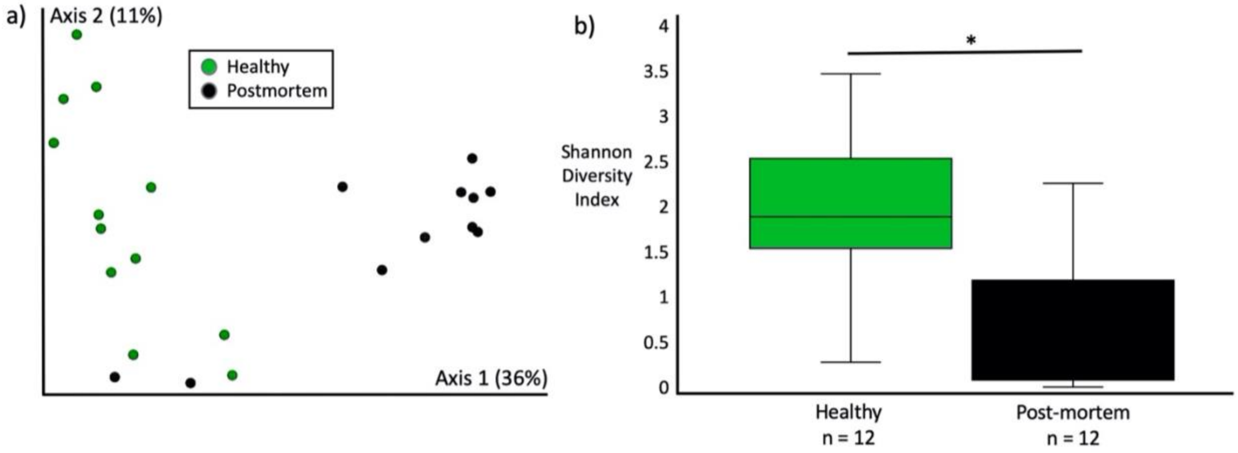

**Additional File 12: Susceptible CZA lorikeets have altered microbial composition that predicts enteritis.** Healthy lorikeets that never developed enteritis throughout the sampling period were identified as “True Healthy” while healthy birds that developed enteritis at least once during the sampling period were identified as “Susceptible.” “Enteritis” represents birds with enteritis that were sampled while they were clinically ill. No post-mortem samples are included in this figure. **a)** Microbial diversity (Shannon, Kruskal-Wallis, $p < 0.005$) was increased in Susceptible and True Healthy birds as compared to birds with enteritis and **b)** microbial composition (Unweighted UniFrac) was altered in Susceptible birds. Variables associated with susceptibility or health were then predicted by a **c)** Random Forest (RF) or **d)** LASSO model. The RF model has a sensitivity of 0.75, a specificity of 0.571, and an overall accuracy of 0.557. This model identifies the relative importance of each variable but not whether the variable is associated with susceptibility or health. The LASSO model has a sensitivity of 0.875, a specificity of 0.571, an overall accuracy of 0.733, and generates an area under the curve (AUC) of 0.72. This model (LASSO) identifies whether a variable is associated with susceptibility or health but not the relative importance of the variable.

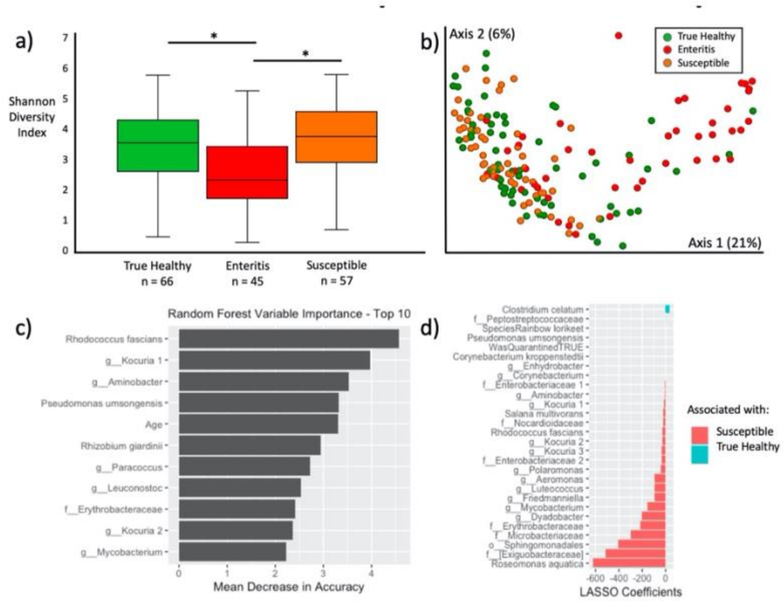
